# 3D chromatin compartment of round spermatids encodes the spatiotemporal program of histone-to-protamine exchange in spermiogenesis

**DOI:** 10.64898/2026.03.10.710708

**Authors:** Mashiat Rabbani, Zachary Apell, Timothy J. Parnell, Lindsay Moritz, Sion Kim, Sowmya Srinivasan, Ritvija Agrawal, Alexander Vargo, Peter Orchard, Wenxin Xie, Lydia Freddolino, Alan P Boyle, Jun Z. Li, Bluma J. Lesch, Bradley Cairns, Minji Kim, Thomas E. Wilson, Saher Sue Hammoud

## Abstract

Sperm formation requires a radical chromatin reorganization, where nucleosomes are replaced by transition proteins (TNPs) and subsequently by protamines (PRM1 and PRM2). Although essential for fertility, the regulatory logic governing this exchange is unknown, but it’s presumed to be stochastic and unregulated. Using endogenously tagged PRM mouse models and stage-resolved, genome-wide profiling, we revise the order of histone-to-protamine exchange and show that the chromatin remodeling process is highly programmed. Imaging and biochemical experiments reveal a direct histone-to-PRM1 exchange, while TNPs appear after PRM1 but precede PRM2 incorporation. This temporal uncoupling of PRM1 and PRM2 incorporation coincides with dynamic, region-specific chromatin remodeling that is not governed by histone acetylation but is instructed by the three-dimensional nuclear architecture of round spermatids. Therefore, by integrating ATAC-seq, CUT&Tag, and Hi-C, we define a compartment-encoded “blueprint” that prescribes the assembly of the mature sperm epigenome and establishes a complex molecular hierarchy for the histone-to-protamine exchange.

## Introduction

In sexually reproducing species, the sperm cell genome undergoes one of the most remarkable chromatin transformations in biology: the transition from a nucleosome-based to a protamine-based chromatin landscape. This process relies on a series of testis-specific basic proteins to achieve the tight chromatin compaction required for sperm function, while preserving the sperm genome’s ability to rapidly and faithfully reactivate after fertilization.^1,2^ The prevailing ’histone-to-protamine’ exchange model depicts this chromatin transformation as a series of stepwise remodeling events in which canonical histones are first replaced by histone variants^3–6^, which overlap with transition proteins (TNP1 and TNP2), before being ultimately replaced by protamines (PRM1 and PRM2).^7–11^ This remodeling cascade implicitly assumes a linear, hierarchical exchange of proteins, in which each intermediate is a prerequisite for the next.

However, accumulating genetic evidence challenges this strict sequence of remodeling events. For example, single *Tnp1* or *Tnp2* knockouts in mice result in sub-fertility^8,10,12^, while double knockouts are male sterile^13^, indicating partial redundancy and cooperative roles in chromatin remodeling.^6, 7^ Notably, PRM1 incorporation occurs normally even in the complete absence of both transition proteins, whereas PRM2 incorporation is compromised, demonstrating that TNPs are required only for PRM2 deposition.^4,6,10^ Likewise, PRM1 deposition is unaffected in *Prm2*-deficient testes, establishing that PRM1 and PRM2 incorporation are mechanistically uncoupled.^8^ Together, these findings point to a modular chromatin-remodeling process, yet how this nuclear repackaging is regulated - and whether it follows a deterministic or stochastic logic - remains unknown.

Despite histone-to-protamine exchange being essential for sperm function, the exchange process is incomplete, with a small subset of histones retained ^14–16^ at developmental gene promoters and at certain repetitive regions of the genome.^17–19^ These retained nucleosomes have been implicated in paternal epigenetic inheritance^14–17,20–22^ and are often presumed to be the only information carriers within the sperm nucleus. This view has prevailed because protamines are presumed to bind DNA uniformly and without sequence specificity. Yet this assumption has not been tested in vivo because of longstanding technical limitations. As a result, whether protamines transmit any information, locus- or compartment-level, to the zygote, or whether they function purely as structural condensation proteins with random genomic distribution, remains an open question.

Here, our findings challenge the established sequence of histone-to-protamine exchange events and the long-standing assumption that PRM1 and 2 are deposited uniformly throughout the paternal genome. By tagging the endogenous *Prm1* and *Prm2* loci with V5 and/or HA epitopes, we demonstrate that PRM1 and PRM2 incorporation is temporally uncoupled, with PRM1 undergoing a direct histone-to-PRM1 transition, whereas PRM2 incorporation occurs only after the appearance of transition proteins. This temporal uncoupling raised the possibility that PRM1 and PRM2 are not uniformly distributed across the genome. To test this, we applied stage-resolved ATAC-seq and CUT&Tag to capture successive waves of chromatin compaction and compositional change, as proxies for PRM1- and PRM2-driven accessibility changes. We hypothesized that if accessibility varies along the genome and across stages this can indicate that protamine exchange is a regionally partitioned process that likely encodes heterogeneity into the sperm chromatin landscape. Our dense, stage-resolved mapping of chromatin accessibility, nucleosome occupancy, and histone modifications indeed confirms a continuous, highly ordered, and progressive change in accessibility across spermiogenesis. This loss of chromatin accessibility coincides with the loss of H2B and H4, indicating that chromatin compaction is coupled to histone eviction. To examine whether H4 hyperacetylation regulates these stage-specific changes, we performed H4ac CUT&Tag across stages and show that H4 acetylation marks early, intermediate, and late remodeling loci-suggesting it accompanies but does not direct the remodeling process. Instead, integrating accessibility dynamics with 3D genome architecture in round spermatids showed that differential accessibility loss across stages is correlated with higher-order genome organization, with euchromatic A-compartment regions compacting earlier than heterochromatic B-compartment regions. Together, these results demonstrate that the temporal uncoupling of PRM1 and PRM2 coincides with a programmed, compartment-encoded remodeling process, suggesting that protamine 1 and -2 incorporation encodes compartment-level architectural information within the mature sperm nucleus.

## Results

### A revised temporal sequence of histone-to-protamine exchange

The precise temporal relationships among histones, transition proteins, and individual protamines in vivo have remained difficult to resolve due to limitations in available reagents and antibodies. To directly define the timing of protamine incorporation during spermiogenesis, and to monitor protamine deposition relative to other nuclear proteins under physiological conditions, we therefore used CRISPR-Cas9-mediated genome editing to introduce V5 or V5–HA epitope tags in frame into mouse *Prm1* and *Prm2* alleles, respectively (Figure 1A; note other tested tags led to infertility). Importantly, both tagged *Prm1* and *Prm2* alleles supported normal spermatogenesis, as evidenced by the normal testis histology, sperm counts, motility, and fertility relative to wild-type controls (Supplementary Figure 1A, B).

**Figure 1:**
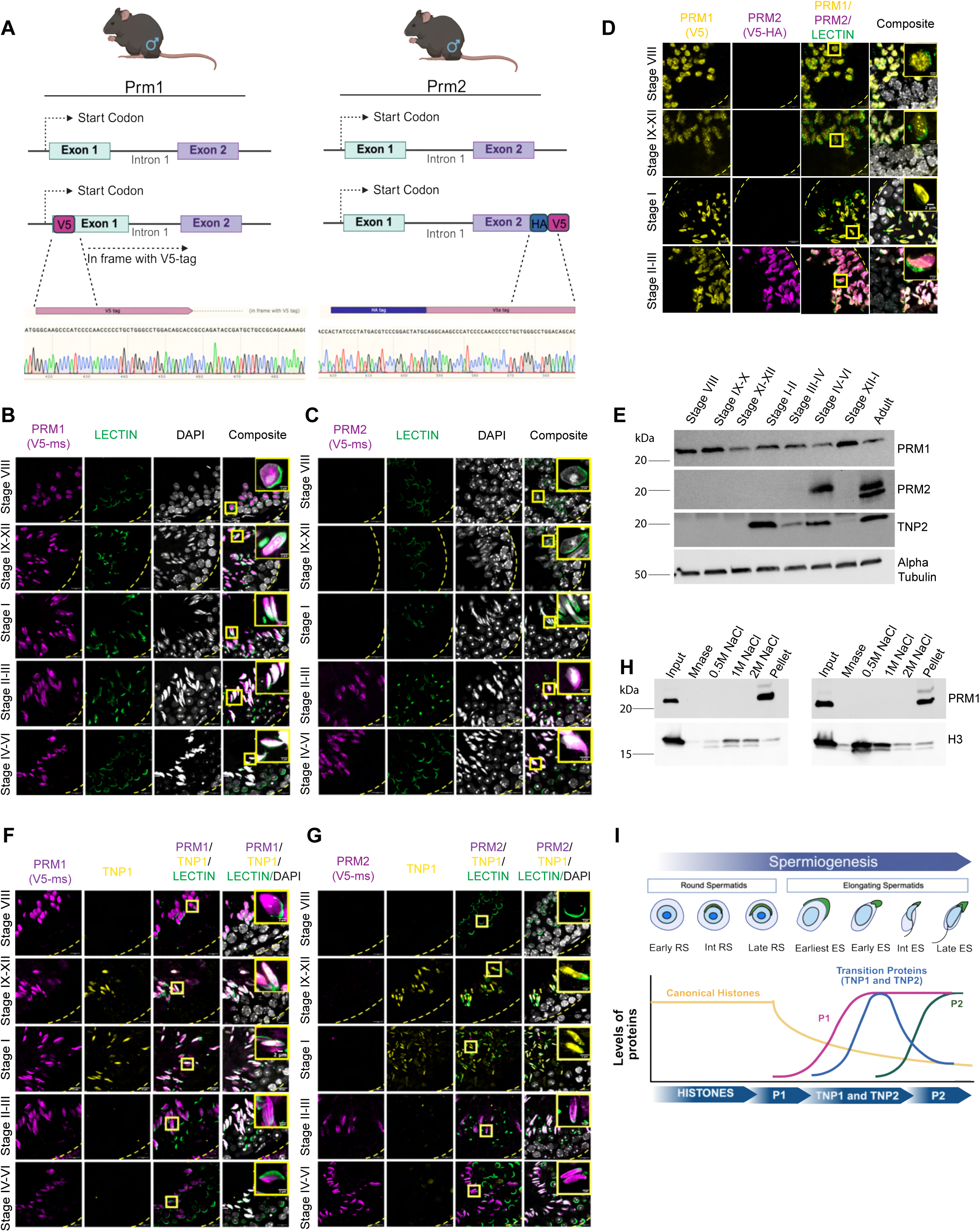
PRM1 undergoes direct histone exchange independently of and prior to PRM2 incorporation during spermiogenesis. **(A)** Schematic of the *Prm1-V5* and *Prm2-HA-V5* knock-in mouse models, with Sanger sequencing confirming in-frame insertion of the V5 tag at the N-terminus of PRM1 and dual HA-V5 tags at the C-terminus of PRM2. **(B)** Immunofluorescence of seminiferous tubule cross-sections from *Prm1-V5* heterozygous mice stained with anti-V5. Roman numerals denote the tubule stage, ordered here according to the sequential progression of histone-to-protamine exchange during spermiogenesis. PRM1 (anti-V5; Magenta); Lectin (green); DAPI (grey). Scale bar: 10 µm; inset: 2 µm. **(C)** As in (B), we co-stained testis cross-sections from *Prm2-V5* heterozygous mice with PRM2 (anti-V5; Magenta), Lectin (green), and DAPI (grey). Scale bar: 10µM, Inset Scale bar 2 µm. **(D)** Co-immunostaining of testes cross-sections spanning stages of seminiferous tubule, stained with anti-V5 and anti-HA in compound heterozygous *Prm1*^V5/+^; *Prm2*^V5-HA/+^ mice. Scale bar: 10µm, Inset Scale bar 2 µm. PRM1 (anti-V5; yellow); PRM2 (anti-HA; magenta), Lectin (green); DAPI (grey). **(E)** Immunoblots of PRM1, PRM2, and TNP2 from whole testes of WIN 18,446-synchronized mice collected at various days post-RA injection corresponding to seminiferous tubule stages VIII, IX–X, XI–XII, I–II, III–IV, V–VI, and IX–X. The adult testis serves as a positive control. **(F)** Co-staining for PRM1 and TNP1. Scale bar: 10 µM, Inset Scale bar 2 µm. PRM1 (magenta); TNP1 (yellow); Lectin (green); DAPI (grey). **(G)** Co-staining for PRM2 and TNP1. Scale bar: 10 µM, Inset Scale bar 2 µm. PRM2 (magenta); TNP1 (yellow); Lectin (green); DAPI (grey). **(H)** Western blot of nuclear lysates from WIN 18,446-synchronized testes isolated at stages IX and X post-RA injection. Chromatin was solubilized by MNase digestion, followed by sequential salt extractions (0.5–2 M NaCl), and immunoblotted for PRM1 and H3 to distinguish chromatin-bound from soluble protamine fractions. **(I)** Schematic for the revised sequence of PRM1 and PRM2 nuclear incorporation during histone-to-protamine exchange, as supported by this figure.

We first analyzed the *Prm1*-V5 and *Prm2*-V5–HA alleles independently to establish their individual timing of nuclear enrichment. Since spermatogonial stem cell differentiation is an asynchronous, highly regulated process, any given tubule cross-section contains germ cells at multiple stages of differentiation. These recurring cellular associations define the 12 stages of the seminiferous epithelium cycle^23,24^ . Within these stages, spermatids mature through 16 morphologically distinct steps. Notably, histone-to-protamine exchange initiates at stage VIII (RS step 8) and proceeds sequentially through stages IX-XII (steps 9-12), I–III (steps 13-14), and IV-VII (steps 15-16). Using either anti-V5 for PRM1 or anti-V5/HA to track PRM2, we observed that PRM1 accumulates in spermatid nuclei at stage VIII and persists through subsequent stages (Figure 1B). In contrast, PRM2 nuclear enrichment was not detected until late stages of spermatid maturation (Stages II-VI; Figure 1C). These observations reveal a previously underappreciated temporal uncoupling between the two protamine proteins (Figure 1B, C). To determine whether this uncoupling reflects bona fide differences in protamine dynamics within the same cellular context - rather than variability across animals or sections - we next examined the relative incorporation of PRM1 and PRM2 within individual cells within a seminiferous tubule. Co-staining of testis sections from *Prm1*-V5; *Prm2*-V5-HA mice with antibodies against V5 and HA revealed the same reproducible temporal pattern observed in the single-allele analyses (Figure 1D). This result was independently confirmed by co-staining *Prm2*-V5–HA testes with an antibody against endogenous PRM1 (Hup1N) (Supplementary Fig. 1C). To further exclude that the observed uncoupling is due to differential nuclear accessibility of the antibodies to DNA-bound PRM1 vs. PRM2, we synchronized spermatid development by modulating retinoic acid signaling.^25^ This approach produces testes in which entire seminiferous tubules are restricted to one to two consecutive stages of the seminiferous cycle (e.g., VII-VIII, IX-X, XI-XII, etc.). Using testis samples containing spermatids at different stages, we performed immunoblots to monitor the dynamics of PRM1 and PRM2. Consistent with the immunofluorescence data, PRM1 protein was detected earlier than PRM2 across synchronized developmental windows (Figure 1E). Together, these complementary approaches establish that PRM1 and PRM2 translation and nuclear localization are temporally uncoupled and differentially regulated.

We next examined the temporal relationship between protamine incorporation and transition protein localization during spermiogenesis. Co-staining of Prm1^v5/+^ testis sections with anti-V5 and either TNP1 (Figure 1F) or TNP2 (Supplementary Figure 1D) revealed that PRM1 accumulates in spermatid nuclei beginning at stage VIII, before detectable nuclear localization of either transition protein, which first appears at stages IX-XII (Figure 1F). This temporal ordering was independently confirmed by immunoblots from the synchronized testes preparations from above, which revealed that PRM1 protein accumulation indeed precedes the appearance of TNPs (Figure 1E). In contrast, co-staining of PRM2^v5/+^ mouse testis sections with anti-V5 and either TNP1 (Figure 1G) or TNP2 (Supplementary Figure 1E) demonstrated that transition proteins precede PRM2 nuclear localization, consistent with the conventional model of transition protein-PRM2 incorporation order.

Although these data show that PRM1 precedes transition proteins, the PRM1 nuclear localization alone does not equate to chromatin association. To determine whether PRM1 in early spermatid is stably incorporated into chromatin or is in the nucleoplasm awaiting transition protein-mediated remodeling, we synchronized spermatogenesis in wild-type mice^25,26^, collected testis enriched for stages VIII or IX of the seminiferous tubule cycle, and subjected single-cell suspensions from these stages to sequential salt fractionation^27^ (Figure 1H). In stage VIII-enriched chromatin, histone H3 was progressively released with increasing salt concentration as expected, whereas PRM1 remained predominantly in the insoluble chromatin pellet even at high ionic strength, indicating stable chromatin association. Notably, the identification of PRM1 in the insoluble fraction is unlikely to be an artifact of aggregation due to disulfide bond formation, as disulfides only form during epididymal maturation^28–30^. This salt-resistant association persisted in stage IX-enriched chromatin, despite increased histone lability at this stage, consistent with the incorporation of testis-specific histone variants^5,6,31^ known to destabilize nucleosomes^32,33^. Together, these results demonstrate that PRM1 is stably chromatin-bound prior to, and independent of, transition protein nuclear accumulation. Consistent with a direct histone-to-PRM1 transition, co-staining of *Prm1*^V5/+^ testes for PRM1, histone H3, or acetylated Histone H4 (H4ac) revealed substantial temporal overlap between PRM1 and Histone H3 or PRM1 and H4ac in stage VIII spermatids (Supplementary Figure 1F,G), but in stage IX-XII the PRM1-high regions lose histone H3 or acetylated histone H4 (H4ac) (Supplementary Figure 1F,G), suggesting that PRM1 deposition coincides with histone H3 or H4ac loss (see CUT&Tag data below).

Together, these data establish a direct histone-to-PRM1 transition in which PRM1 is stably incorporated into chromatin prior to the accumulation of transition proteins and independently of PRM2. In contrast, PRM2 incorporation follows transition proteins, as previously described (Figure 1I). We therefore propose a revised sequence for the histone-to-protamine exchange: Histones → PRM1 → TNP1/2 → PRM2 (Figure 1I). This revised sequence of chromatin remodeling is strongly supported by earlier genetic data showing that PRM1 nuclear incorporation is preserved in *Tnp1/Tnp2* double-knockout testes, despite the complete absence of transition proteins, whereas PRM2 incorporation is compromised or lost in some spermatids ^13^, indicating that PRM1 and PRM2 likely incorporate into chromatin through distinct mechanisms. More importantly, this direct histone-to-PRM1 exchange is not an idiosyncrasy of mouse spermiogenesis but has been reported in birds and certain fish, suggesting this is a likely conserved feature of vertebrate spermiogenesis^34,35^.

### Defining stage-specific architectural changes in sperm chromatin during spermiogenesis

The temporal uncoupling of PRM1 and PRM2 incorporation raises the possibility that protamine deposition follows a staged, ordered compaction program rather than a single, uniform compaction event. A definitive test of this model would require mapping PRM1- and PRM2-bound chromatin at high resolution in sperm. However, the extreme compaction of protamine-bound DNA and the lack of accessible linker regions between protamine molecules in sperm make direct nuclease- or transposase-based foot-printing approaches technically impossible^36^. We therefore adopted an indirect but powerful strategy, using ATAC-seq^37,38^ to monitor chromatin accessibility dynamics across spermatid maturation.

To collect spermatids at specific stages of development, we combined spermatogenesis synchronization ^25^ with fluorescence-activated cell sorting^39^ (Figure 2A) to enrich for a homogeneous pool of spermatids isolated from any given stage (Figure 2B; Supplementary Table 1; **see Methods**). Importantly, because elongating spermatids are largely transcriptionally quiescent, the stage-specific changes in accessibility primarily reflect structural changes in chromatin rather than transcriptional activity.^40,41^ Therefore, the accessibility changes we observe can serve as a sensitive proxy for distinct compaction states dominated by histones, PRM1, or PRM1 plus PRM2 (Figure 2A).

**Figure 2:**
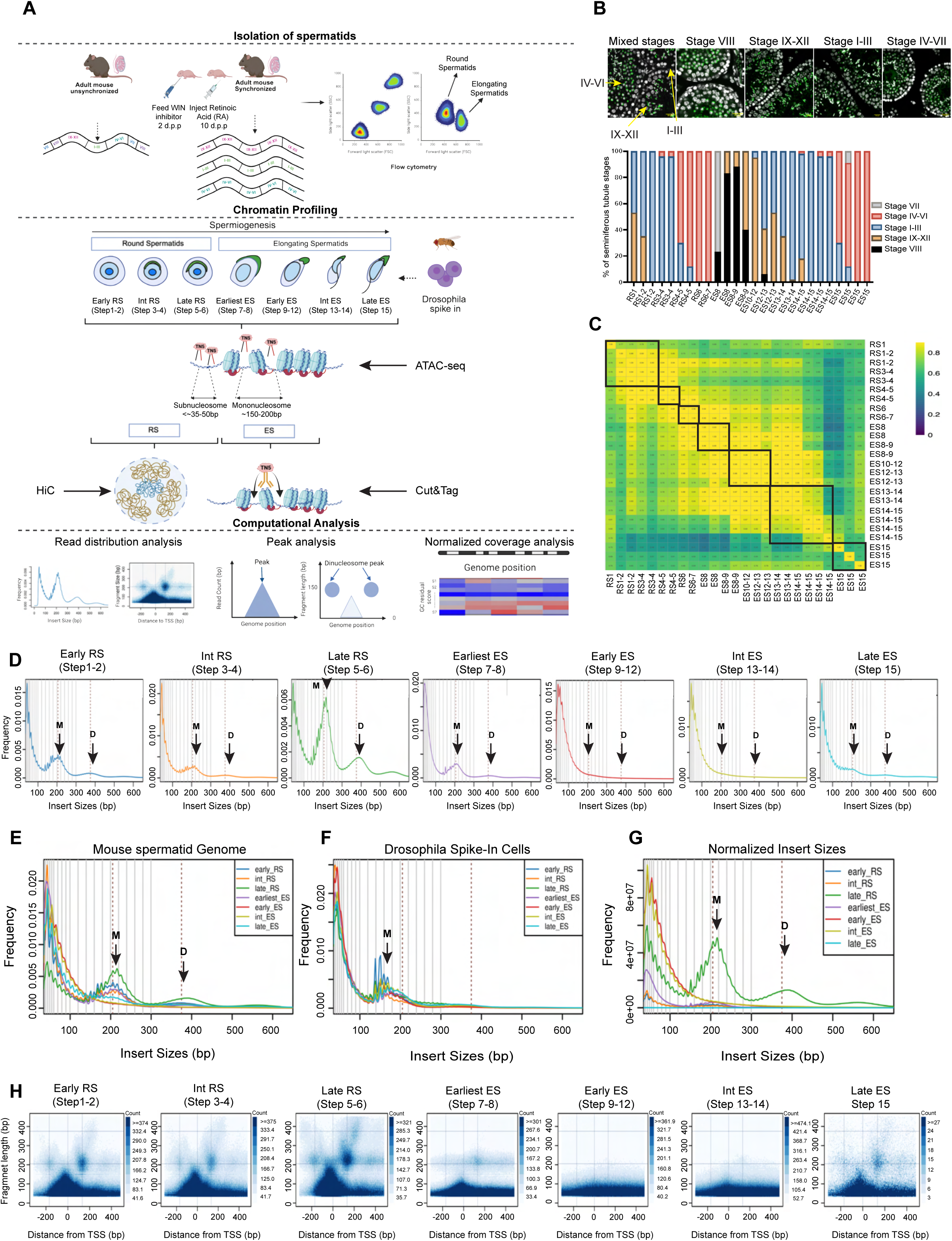
Stage-resolved chromatin profiling reveals progressive and programmatic loss of nucleosome footprint during spermiogenesis. **(A)** Schematic of the experimental design and computational framework for stage-specific chromatin profiling across spermiogenesis. Mouse spermatid populations were isolated from specific stages of the seminiferous tubule cycle by flow cytometry from unsynchronized and WIN 18,446/RA-treated testis and subjected to ATAC-seq, CUT&Tag, and Hi-C. Computational analyses included read distribution analysis, base-level peak calling, and GC-normalized bin-level coverage analysis. **(B)** Representative immunofluorescence images of seminiferous tubule cross-sections from unsynchronized and synchronized mice (top). Bar plot (bottom) shows the proportion of tubules at each seminiferous tubule stage across collected samples, confirming stage enrichment by synchronization. **(C)** Spearman correlation heatmap of all ATAC-seq samples. Biological replicates cluster tightly and samples separate by developmental stage. **(D)** ATAC-seq fragment length, i.e., insert size frequency distributions aggregated across the seven collapsed datasets of spermiogenesis. Solid vertical lines are size boundaries used during data analysis (see Methods). Arrows at 205 and 375 bp label the centers of the mononucleosome (M) and dinucleosome **(D)** fragment peaks in each stage. **(E)** Overlaid ATAC-seq fragment length distributions from early round to elongating spermatid populations. Arrows at 205 and 375 bp label the centers of the mononucleosome (M) and dinucleosome (D) fragment peaks. **(F)** Overlaid ATAC-seq fragment length distributions from *Drosophila* spike-in reads across all seven spermatid stages, for comparison to (E). Arrow at 165bp labels the center of the mononucleosome (M) fragment peak. **(G)** Overlaid fragment length distributions from (D) after normalizing to the corresponding *Drosophila* spike-in, to reveal changes in absolute chromatin accessibility between stages. Arrows at 205 and 375 bp label the centers of the mononucleosome (M) and dinucleosome (D) fragment peaks. **(H)** V-plots of ATAC-seq fragment lengths across all seven stages as a function of distance from the TSS of actively transcribed genes based on PRO-seq (Kaye et al., 2024). The characteristic promoter nucleosome pattern in round spermatids is progressively lost in elongating spermatid stages, reflecting stage-dependent disruption of promoter-proximal nucleosome positioning.

To map chromatin accessibility dynamics over time, we densely sampled spermatid maturation, generating 26 ATAC-seq datasets spanning the 16 steps of spermatid maturation (Figure 2B; Supplementary Table 1). Each sample contained approximately 220,000 mouse cells supplemented with a 10% *Drosophila* spike-in to normalize across samples and to control for transposase activity and library complexity (Figure 2A). To assess reproducibility and global chromatin similarity across spermatid populations, we computed pairwise Spearman correlations of normalized signals across 10kb bins for all 26 datasets (Figure 2C). The resulting clustered heatmap revealed high concordance between biological replicates, which clustered tightly along the diagonal, confirming the robustness of our cell synchronization and sorting. Notably, correlations decreased progressively between non-adjacent stages, reflecting the continuous remodeling of the chromatin landscape during spermiogenesis (Figure 2C). Based on the correlation structure and confirmed biological sample stages (Figure 2B;2C; Supplemental Table 1), we binned samples into seven discrete windows of spermatid maturation (Figure 2A;2B; Supplementary Table 1) spanning early round spermatids (Early RS) through late elongating spermatids (Late ES; isolated from stages I-XII; Steps 1–15). These windows allow us to capture relevant developmental transitions while maintaining the granularity of the spermatid maturation trajectory. The latest elongating spermatid samples we could generate sufficient-quality ATAC-seq libraries without resorting to harsh chemical treatments, such as DTT, to increase chromatin accessibility are Step 15 spermatids isolated from Stage IV of the seminiferous tubule cycle.

By leveraging intrinsic ATAC-seq features, such as insert size distributions at specific genomic loci, such as transcriptional start sites (TSSs), or across the genome, the data provide a global, model-independent readout of the changing spermatid chromatin architecture. Early round spermatids displayed canonical nucleosome footprints, with prominent peaks at approximately 200 bp and 400 bp corresponding to mono- and di-nucleosome footprints (Figure 2D, Figure 2F; Supplementary Fig. 2A). This organization remained stable throughout RS stages, indicating an intact nucleosome organization during this phase of development. However, as cells progressed to Early ES, the nucleosome-associated peaks were lost, and wider subnucleosomal peaks emerged (Figure 2D, 2E, Supplementary Figure 2A). These fragment sizes resemble destabilized nucleosome intermediates observed during active chromatin remodeling in other systems^42–44^. Curiously, these expanded sub-nucleosome peaks persisted through intermediate elongating spermatids (Int ES) but were lost in Late ES (Figure 2F). Concomitant with this loss, we observed a reemergence of mono-nucleosome-sized fragments in Late ES, suggesting re-stabilization of chromatin and selective retention of histones in a very small fraction of the mature sperm genome (Fig. 2D, Supplementary Fig. 2A).

To determine whether the loss of the nucleosome footprint in the early and Int ES was a biological state or a technical artifact, we leveraged the *Drosophila* spike-in data, which shows a well-defined nucleosome footprint across all samples (Figure 2B; Supplementary Figure 2B). Furthermore, when overlaying insert size distributions from all *Drosophila* spike-in datasets, there was no broadening or distortion of nucleosome peaks (Figure 2F), confirming that the loss of nucleosome footprint in ES isn’t due to excessive enzymatic digestion in some samples but a true biological chromatin state.

Having established this, we next asked whether it reflects a globally hyper-accessible chromatin state, perhaps coinciding with the previously described H4 hyperacetylated chromatin state known to loosen chromatin and facilitate histone-to-protamine exchange^45^. To estimate absolute accessibility, we normalized spermatid ATAC-seq signal to that of the *Drosophila* spike-in, which revealed that early and Int ES had markedly increased chromatin accessibility relative to other stages (Figure 2G). Notably, late RS displayed a similarly elevated accessibility relative to spike-in despite retaining canonical nucleosome organization, identifying a hyper-accessible yet nucleosome-preserved intermediate chromatin state (Figure 2G).

To determine how global changes in chromatin accessibility impact the organization of positioned nucleosomes, we generated insert size-resolved V-plots centered on annotated transcription start sites (TSS, ±200 bp), where assignment of genes to transcribed/active and non-transcribed/inactive states was based on previously published PRO-seq data from pooled round spermatids^46^. In RS (early through late), V-plots revealed a canonical promoter architecture characterized by well-positioned +1 and -1 nucleosomes flanking active TSSs (Figure 2H). As cells transitioned from late RS to the ES stage (Stage VIII), we first observed an attenuation of the di-nucleosome signal (earliest ES), followed by a complete loss of positioned nucleosomes (early & Int ES) (Figure 2H). Interestingly, a small fraction of positioned +1 nucleosomes re-emerges in Late ES, a trend that mirrors our genome-wide fragment length analysis (Figure 2D). In contrast, inactive gene promoters in RS lacked a defined nucleosome-free region (NFR) or structured promoters but had a diffuse nucleosome distribution pattern across the promoters, consistent with a transcriptionally silent architecture (Supplementary Figure 2C). Curiously, this disappearance of nucleosome-protected fragments was not restricted to active genes; as both transcribed and inactive promoters displayed a loss of a nucleosome footprint in the RS-to-ES transition, which likely reflects global hyper accessibility and destabilization of the chromatin landscape (Figure 2D, 2H).

Together, our time-resolved ATAC-seq analysis reveals multiple distinct states: a nucleosome-stable state in early and intermediate RS; a transient, hyper-accessible but nucleosome-stable state in late RS and the earliest ES stages; and a hyper-accessible and nucleosome-depleted phase during early-to-intermediate ES. In Late ES, a small but reproducible mono-nucleosome-sized fragment population re-emerges, consistent with partial chromatin re-stabilization and selective histone retention rather than de novo nucleosome assembly (**see below H3K27me3**).

### The spatiotemporal landscape of chromatin remodeling reveals asynchronous compaction of the spermatid genome

To determine whether chromatin compaction in maturing spermatids follows a programmatic process, we analyzed chromatin accessibility changes within and across developmental stages at lower resolution. Because ATAC-seq can be influenced by GC content, and because chromatin composition changes markedly during spermatid maturation, we first examined GC-dependent biases across samples as this can confound comparisons across spermiogenesis (see **Methods**). When examining the ATAC-seq signal as a function of GC content across 1-kb genomic bins, we observed a progressive increase in GC-read count correlation from Early ES to Int ES (Supplemental Figure 2D). This pattern likely reflects the unusually accessible chromatin state at these stages, where increased enzyme accessibility amplifies intrinsic sequence preferences, leading to the exaggerated GC bias. To prevent these stage-specific biases from dominating inter-sample differences, ATAC-seq signal in 1kb bins were regressed as a function of GC content and converted to GC-normalized residual Z-scores. This normalization minimizes sequence-driven effects and facilitates direct comparison of relative chromatin accessibility across stages. A representative GC-normalized ∼1 Mb region on chromosome 9 encompasses multiple genes and distinct patterns (Figure 3A; high Z-scores in red denote more accessible spans, while lower Z-scores in blue denote less accessible spans). Even within this small genomic interval, chromatin accessibility changes vary across loci and developmental stages. For example, Region I overlaps with the *Scaper* locus, which is transcribed in RS (Pro-seq track; red) and shows high accessibility early but progressively diminishes as spermatids elongate. In contrast, Region II, encompassing the non-transcribed gene *Lingo1,* is relatively inaccessible in round spermatids but transiently gains accessibility during early and Int-ES stages, when germ cells are transcriptionally quiescent, and returns to an inaccessible state in Late ES. Region III overlapping gene *Clk3* is highly transcribed in RS and remains accessible across all stages of spermatid development, including Late ES. Together, these representative patterns recur broadly across the genome and demonstrate that chromatin compaction during spermiogenesis proceeds in a region-specific and developmentally ordered manner rather than through uniform genome-wide silencing.

**Figure 3:**
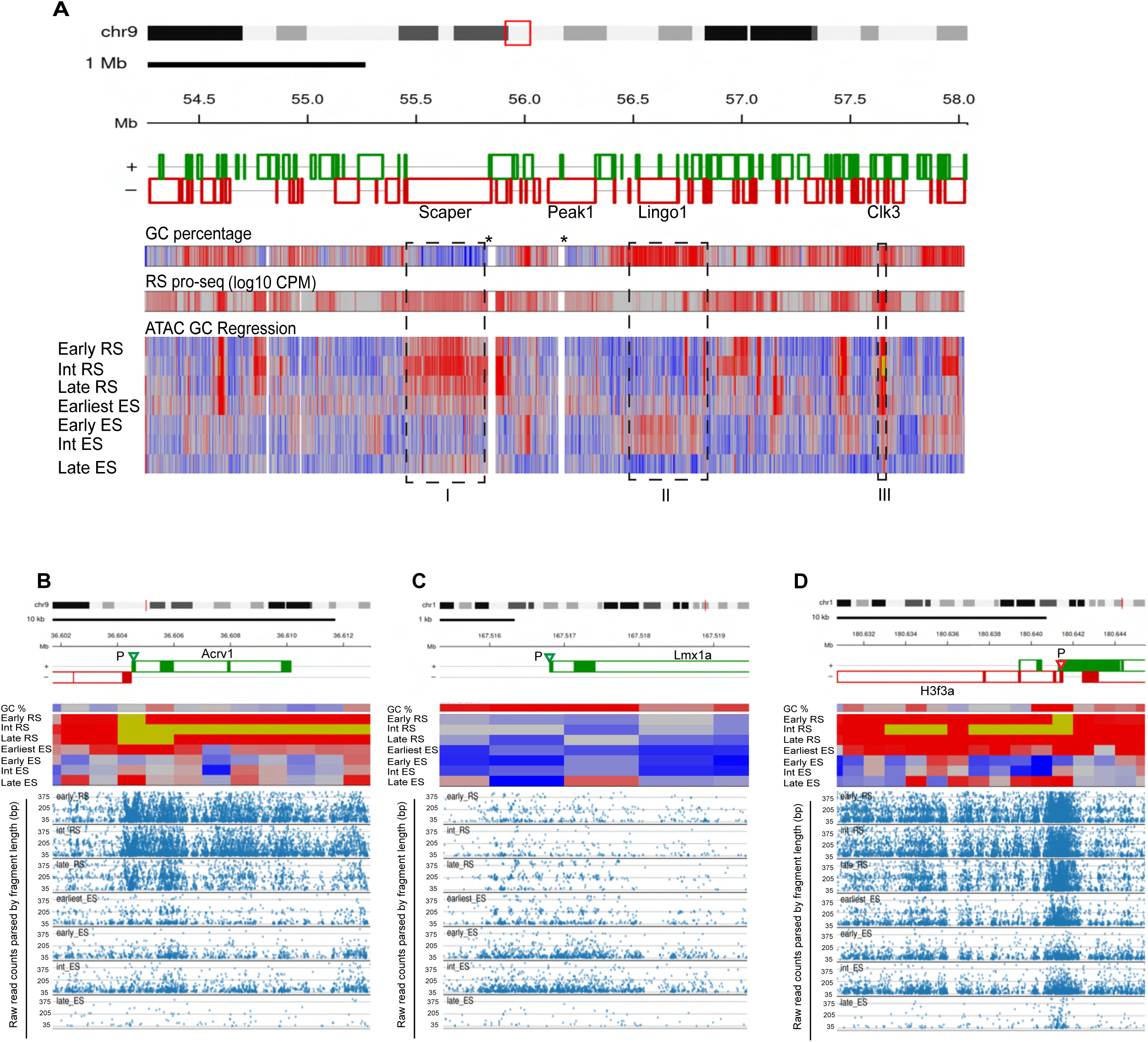
Genome-wide chromatin profiling reveals programmed, locus-specific remodeling during histone-to-protamine exchange. **(A)** Corrected ATAC-seq accessibility across a 4 Mb region of chromosome 9 for all seven spermatid stages (Early RS through Late ES). Tracks from top to bottom: GC content (red, GC higher than the grey genome median; blue, GC lower than median); PRO-seq signal from round spermatids (log₁₀ CPM; red, transcribed) ^30^; GC-residual ATAC-seq Z-scores per stage (red, relatively more accessible than the grey regression median, i.e., Z>0; blue, relatively less accessible, Z<0). Dashed boxes highlight three representative classes of locus behavior: progressive loss of accessibility in elongating spermatids (I); gain of accessibility in elongating spermatids (II); and stable accessibility throughout spermiogenesis (III). Asterisk (*) indicates blacklisted regions containing repetitive elements in the genome. **(B-D)** GC-residual ATAC-seq Z-scores (upper panels like (A); yellow, saturated high Z) and genomic V-plots across all seven stages (lower panels), for three loci exemplifying distinct accessibility trajectories. Each dot is a single mapped insert. Horizontal lines are at 35bp (minimum allowed fragment size), 205 bp (mono-nucleosome peak center), and 375 bp (di-nucleosome peak center). **(B)** *Acrv1* locus with high accessibility in round spermatids that is progressively lost during elongation. **(C)** *Lmx1a* locus with peak accessibility during the Late RS-to-Earliest ES transition. **(D)** *H3f3a* locus that maintains accessibility throughout spermiogenesis, with nucleosome occupancy selectively preserved at the promoter in Late ES.

To examine local chromatin changes during spermiogenesis at higher resolution, we next generated genome-wide V-plots (Figure 3B–D; see **Methods**) to provide a read-level view of nucleosome composition or its disassembly intermediates across spermatid maturation at all loci. At the *Acrv1* locus, a highly transcribed acrosomal protein gene (Supplemental Figure 3A)^15^, V-plots revealed high accessibility in early stages (Early, In, Late RS; yellow heatmap is saturated high accessibility) followed by progressive loss of accessibility during elongation, consistent with transcription-dependent chromatin compaction (Figure 3B). However, the temporal ordering of *Acrv1* remodeling is graded. Nucleosome-sized fragments are first lost from the gene body and are only later lost from the promoter. In contrast, *Lmx1a*, a neuronal gene not transcribed in the male germline (Supplemental Figure 3B), follows a distinct chromatin remodeling trajectory (Figure 3C). Accessibility over the promoter region is diffuse and weakly structured in early and intermediate RS but becomes more pronounced and punctuated in late RS and earliest ES. By early and intermediate ES, the well-defined mono-nucleosome-sized fragments are lost, indicating a delayed, transcription-independent remodeling process (Figure 3C). At the ubiquitously expressed histone H3.3 gene, chromatin accessibility is maintained up to the Late ES stage, at which time gene body coverage is lost but promoter-centered nucleosome fragments are retained (Figure 3D; Supplementary Figure 3C). Finally, to confirm that the observed locus-specific patterns were not influenced by global differences in accessibility or sequencing depth across stages, we normalized insert-size counts to whole-genome signal within each developmental stage, which preserved all qualitative patterns (Supplementary Figure 3D; note samples sequenced to comparable depth). These examples show how chromatin remodeling during spermiogenesis proceeds through multiple, locus-specific patterns of chromatin accessibility gains and losses, reflecting both transcription-dependent and transcription-independent mechanisms. The switch of highest accessibility from region to region across spermiogenesis argues for a genome-scale, programmatic reorganization of chromatin rather than a uniform or stochastic compaction process.

### Peak-level chromatin dynamics support a continuous and programmed genome-wide remodeling during spermiogenesis

To further understand the unique properties of ATAC-seq read re-distribution during spermiogenesis, we next performed peak calling using two independent methods: (i) MACS2^47^ to identify regions of enriched accessibility above the genomic background (Figure 4A), and (ii) a dinucleosome-specific peak-calling algorithm that locates significantly accessible loci with multiple, well-positioned nucleosomes (Figure 4B, **see Methods**). The number of peaks identified by either method deviated significantly from random expectation (Figure 4A, global p=2.84e-11; Figure 4B, global p=8.46e-15). As expected, MACS2 identified more peaks, with the dinucleosome-defined peaks being entirely nested within the MACS2 set in every stage (Supplemental Figure 4A). Thus, while many regions are accessible, only a subset in each stage have the architectural stability of a multi-nucleosomal array. Despite the global chromatin remodeling and loss of nucleosome-sized fragments and positioned nucleosomes during spermatid elongation, the MACS2 peak number increased in early and intermediate elongating spermatids (Early ES: n=364k, p<0.001; Int ES: n=483k, p<0.001) before decreasing greatly in late elongating spermatids (Late ES: n=62k, p<0.001; Figure 4A). In contrast, regions maintaining stable dinucleosomes decreased the most in the Early ES and Int ES stages (Figure 4B). This inverse relationship between MACS and the dinucleosome algorithm indicates on a genome-wide scale that while Early ES and Int ES cells have highly accessible genomes, this accessibility lacks a stereotypical nucleosomal signature (Figure 2D, 2E, 4A, 4B). Notably, both methods identified a significant number of accessible loci in late ES, with peaks from each enriched at promoters relative to the genome average, especially the dinucleosome-enriched peaks (Figure 4C, D). When comparing dinucleosome accessible regions between RS and Late ES, we find that 6118 out of 6219 (98.4%) Late ES peaks overlapped a peak location seen in round spermatids, indicating that they do not represent acquired chromatin pattern during spermatid maturation but are a subset of early RS peaks that reemerge once the period of extreme accessibility resolves. Although round spermatid contamination can never be 100% ascertained, several lines of evidence argue against this interpretation: 1) we monitored purity of cells after FACs sorting, we pre-treated samples with DNase during sample preparation (reducing the possible of cell free chromatin), 3) when comparing global differences in chromatin accessibility patterns outside promoters using MACs peaks ∼50% of loci overlap suggesting that these samples as whole differ from RS. Consistent with this, the B-compartment regions are only accessible in ES samples, not in RS (see below). Furthermore, gene ontology (GO) analysis for enriched promoters in all RS, early, and Int ES tended to enrich for the spermatogenesis program, DNA repair, and cilium formation (Supplementary Table 2), consistent with the expected developmental programming of these cells. In Late ES, GO recovered terms with well-established roles during embryonic development, including chromatin-remodeling, DNA repair, and developmental processes (Supplementary Table 2).

**Figure 4:**
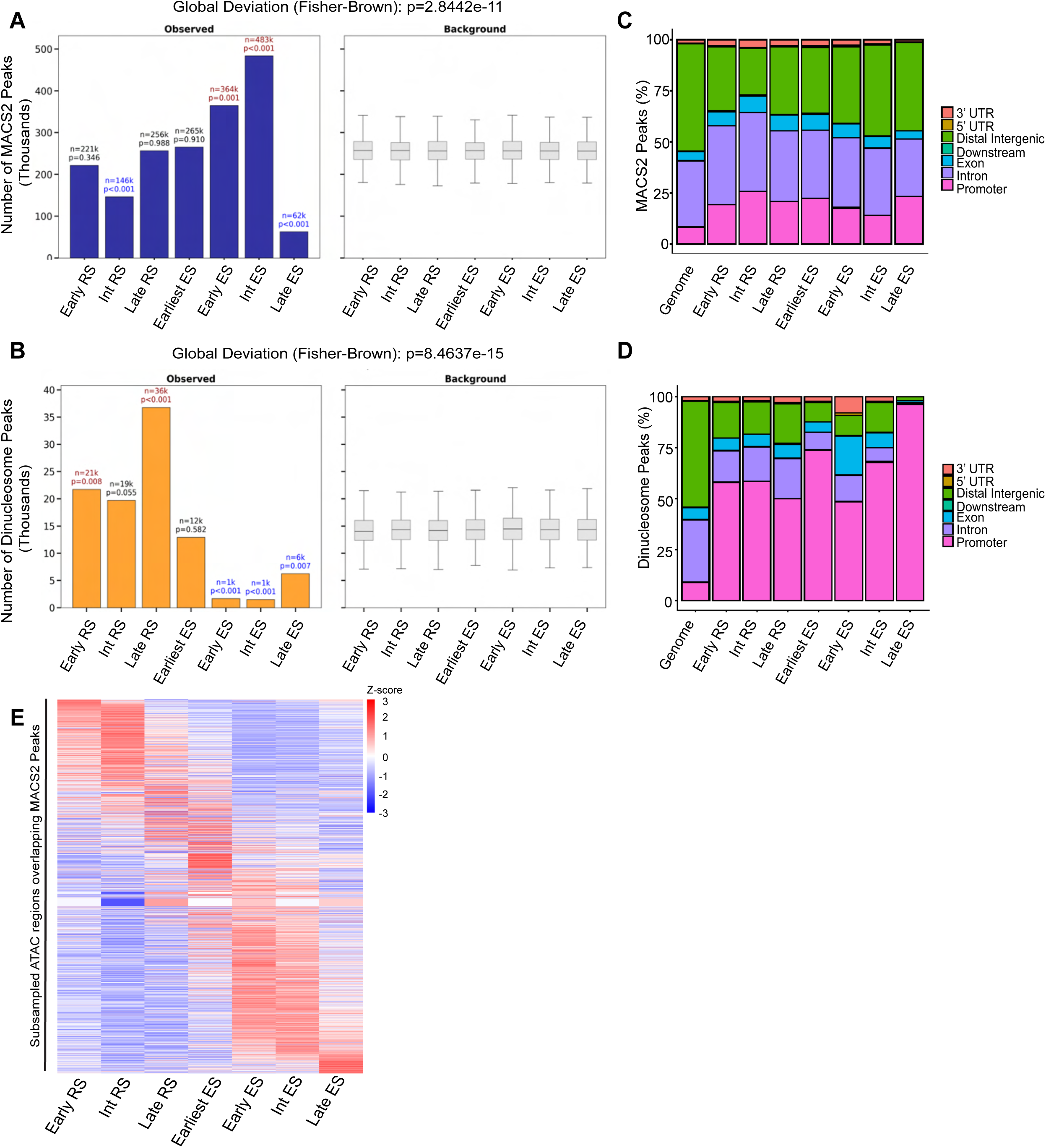
Genome-wide peak analysis confirms a programmed and continuous chromatin remodeling process, with selective retention of intact nucleosomes at promoters. **(A)** Bar plot of the number of accessible peaks identified by MACS2 at each spermatid stage (left), with observed peak counts annotated with Benjamini-Hochberg-adjusted p-values relative to a stage-matched null distribution (right). The global p-value (Fisher-Brown method, p=2.84×10⁻¹¹) indicates that the overall distribution of peak counts deviates significantly from random expectation across spermiogenesis. **(B)** As in (A), for peaks identified by the dinucleosome-specific peak-calling pipeline (global Fisher-Brown p=8.46×10⁻¹⁵). **(C)** Genomic distribution of MACS2-called peaks across spermatid stages, expressed as proportion of peaks overlapping annotated genomic features. **(D)** Genomic distribution of dinucleosome-defined peaks across spermatid stages, as in (C). Both peak calling methods show enrichment at promoters relative to the genome average. **(E)** Heatmap of GC-regression-corrected ATAC-seq Z-scores at subsampled MACS2-defined peaks called in at least one stage. Each row is a subsampled peak overlap region ordered by the centroid of the stage-specific signals.

Finally, to characterize the temporal dynamics of ATAC-seq peaks, we either aggregated all MACS or all dinucleosome-defined peaks across stages and quantified their accessibility profiles over spermatid maturation. Consistent with the previous GC-normalized genome-wide patterns, these peaks display clear locus- and stage-dependent changes in chromatin accessibility. (Figure 4E; Supplementary Figure 4B). Principal component (Supplementary Figure 4C) and UMAP analysis (Supplementary Figure 4D) of peak-level accessibility reveal a continuous trajectory that parallels the temporal progression of spermiogenesis, with biological replicates clustered tightly and adjacent stages occupying neighboring positions along the trajectory. Together, these analyses indicate that the chromatin transition during spermiogenesis, even at peaks, occurs through a gradual, highly coordinated process, rather than a stochastic process.

### Chromatin remodeling in elongating spermatids is decoupled from transcription, driven by targeted histone eviction, and not directed by H4 acetylation

To understand whether the accessibility changes observed between late round/earliest ES (spermatids in stages 7-8) and Early ES (spermatids in stage 9-12) are transcription-dependent, we integrated our GC-normalized ATAC-seq profiles with stage-matched scRNA-seq centroids^48^ and quantified the coupling between promoter accessibility and gene expression across spermatogenic stages. Across all round spermatid stages, promoter accessibility and transcription dynamics are tightly coupled (Figure 5A, Supplementary Figure 5A). In contrast, this coupling between chromatin accessibility and transcription was lost in Early ES (Figure 5A, Supplementary Figure 5A); consistent with the known transcriptional quiescence in elongating spermatids. Interestingly, genes in the top 5% of transcriptional activity in early RS not only show a steeper decline in accessibility during elongation than moderately transcribed genes (25–50th percentile), but by Early ES they become less accessible than regions that were previously lowly transcribed - a reversal that cannot be explained simply by their higher starting accessibility (Figure 5B). This suggests that highly transcribed genes are preferential substrates for chromatin remodeling during the RS-to-ES transition, and that prior transcriptional activity may license or direct early compaction.

**Figure 5.**
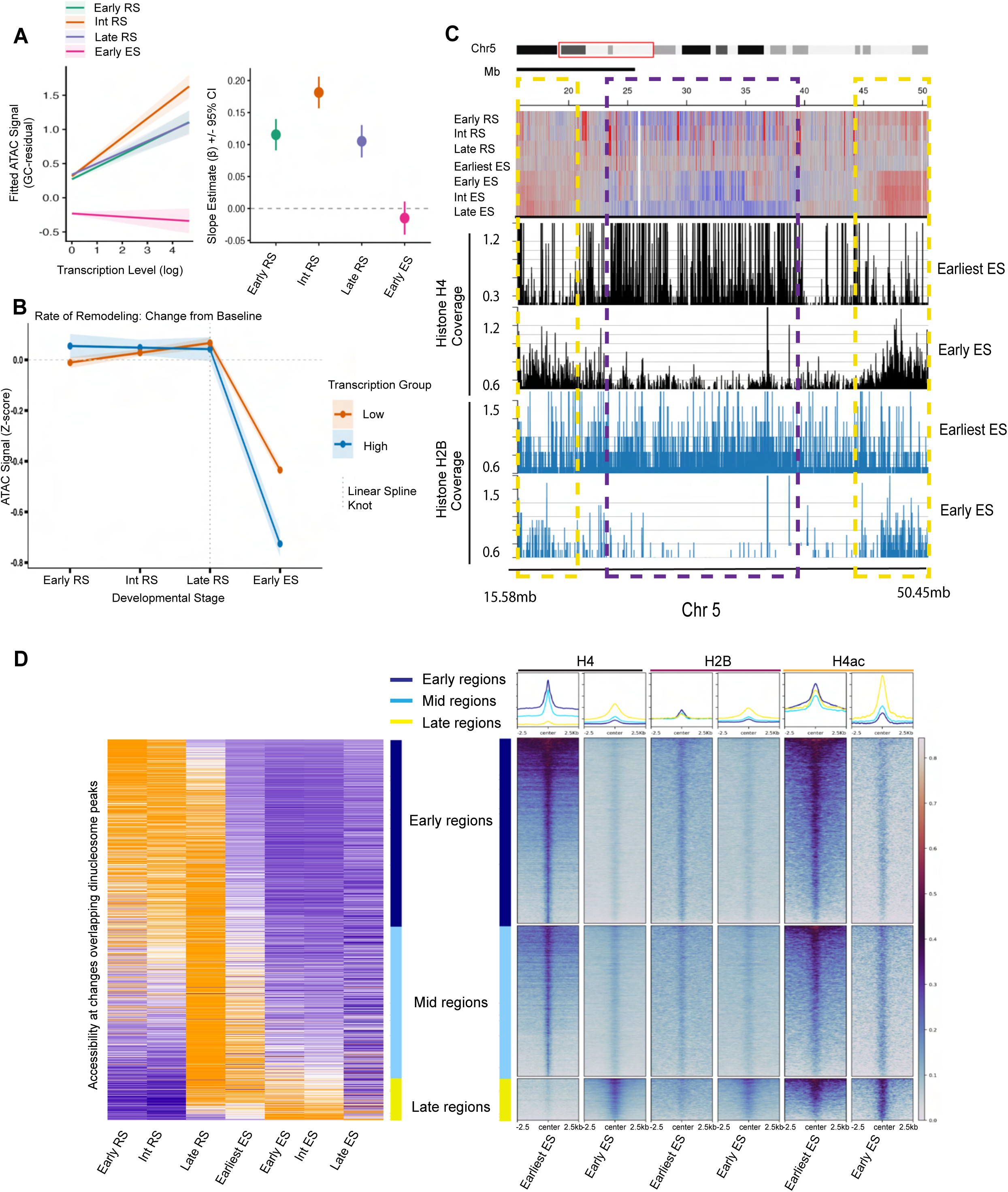
Chromatin accessibility loss during spermiogenesis is coupled to RS transcriptional activity and coincides with direct histone eviction. **(A)** Relationship between transcription level and chromatin accessibility across spermatid stages. Left: marginal predictions from a linear mixed-effects model showing fitted ATAC-seq signal (GC-corrected) as a function of transcription level (log₁₀ CPM) at Early RS, Int RS, Late RS, and Early ES. Shaded ribbons denote 95% confidence intervals. The positive transcription-accessibility coupling present throughout RS stages (green, orange and purple) collapses to zero in Early ES (magenta). Right: stage-specific slopes (β) with 95% confidence intervals. **(B)** Model-estimating change in chromatin accessibility during spermatid development (GC-residual ATAC-seq Z-score) for genes in the top 5% of transcriptional activity (High; blue) and the 25–50th percentile (Low; orange) in Early RS. **(C)** GC-regression-corrected ATAC-seq Z-scores across a representative region of chromosome 5 (top), with CUT&Tag read coverage for histone H4 (black) and H2B (blue) at Earliest ES and Early ES shown below. Regions enclosed by purple dashed boxes or yellow dashed boxes exemplify differences in accessibility and histone eviction. **(D)** (left) Heatmap of GC-residual ATAC-seq Z-scores at dinucleosome-defined peaks. Each row is a subsampled peak overlap region ordered by the centroid of the stage-specific signals. Assigned groups based on dinucleosome peak calls reflect the temporal remodeling class: Early (accessibility lost first), Mid, and Late. (right) CUT& Tag meta-gene heatmaps for H4, H2B, and H4ac over Early, Mid, and Late remodeling regions at Earliest ES and Early ES.

Given the loss of transcription-accessibility coupling in ES, we hypothesized that the regions that gain and lose accessibility are likely due to changes in chromatin composition rather than changes in transcription. This interpretation is consistent with the observed onset of PRM1 incorporation during very late rounds and early elongating (Figure 1B). To test whether accessibility loss corresponds with histone eviction, we profiled histone H4 and H2B by CUT&Tag in late rounds/earliest elongating or early elongating spermatids and compared histone abundance at early-, mid-, and late remodeling loci defined from our ATAC-seq dataset. Regions that lose relative accessibility from late rounds/earliest ES to Early ES also lose both H2B and H4 signal as compared to adjacent regions (Figure 5C). This pattern is not specific to a representative locus on chromosome 5 but is seen at all early- and mid-remodeling loci (Figure 5D), suggesting that accessibility loss in Early ES is likely due to nucleosome eviction and transition to a less accessible epigenomic state. To determine what might guide this selective histone eviction, we examined histone H4 acetylation - a modification associated with chromatin loosening and proposed to facilitate histone-protamine exchange during spermatid remodeling across multiple species^45,49–51^. CUT&Tag profiling of H4ac in late RS/earliest ES shows that H4ac, like H4 and H2B, is broadly distributed across all groups, with comparable enrichment in regions regardless of whether regions remodel early or late (Figure 5D; right panel). In Early ES, H4ac is markedly reduced in early-and mid-remodeling regions but is selectively retained at late-remodeling loci. Thus, H4ac marks chromatin competent for eviction but does not specify the temporal order of eviction, implying that additional mechanisms dictate the spatiotemporal hierarchy of chromatin remodeling (Figure 5D).

### Large-scale programmatic closing of the spermatid genome is defined by A and B compartments

Although individual loci display stage-specific accessibility changes during spermiogenesis (Figure 3), chromatin changes appear to be coordinated over local contexts. To quantify this coordination, we performed a variogram analysis, which showed that variance increases with inter-peak distance up to a plateau of maximal inter-peak variance at ∼100 kb (Supplementary Figure 6A), indicating that nearby regions tend to remodel together over a characteristic neighborhood-scale length. To test whether nuclear organization of chromatin may facilitate this coordination, we generated four Hi-C datasets from round spermatids (day 35 and day 39 post-synchronization), prior to the onset of histone-to-protamine exchange, to delineate round spermatid-specific A/B compartments. Biological replicates showed high concordance (HiCRep stratum-adjusted correlation coefficients of ∼0.83–0.92; Supplementary Figure 6B), providing justification to combine and create a single high-resolution round-spermatid Hi-C dataset totaling ∼2.3 billion alignable reads (**see Methods**). As expected, the resulting Hi-C contact maps revealed strong A/B compartmentalization and preferential self-interaction within compartments (Figure 6A). We confirmed that the defined compartments have the predicted genomic features, e.g., A-compartment regions are enriched for higher GC content (%GC track) and transcriptional output (PRO-seq track), whereas B-compartment regions displayed the opposite pattern (Figure 6B).

**Figure 6:**
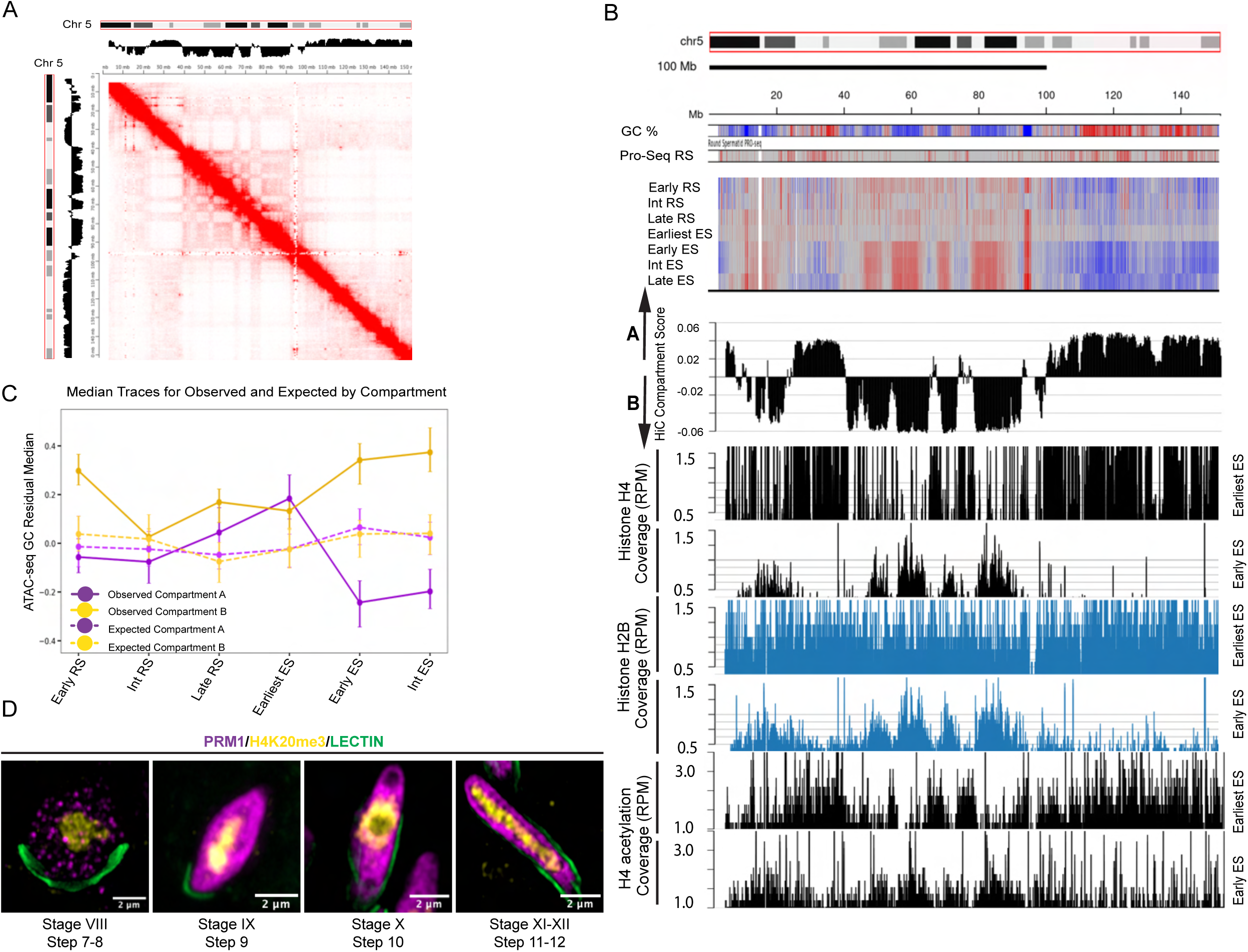
Chromatin compartment identity governs the spatial and temporal order of histone-to-protamine exchange. **(A)** Hi-C contact matrix of intrachromosomal interactions on chromosome 5 from round spermatids, with the corresponding A/B compartment track shown above and to the side (positive values, A compartment; negative values, B compartment). **(B)** Track view of chromosome 5 showing, from top to bottom: GC content per bin; PRO-seq signal from round spermatids; GC-regression-corrected ATAC-seq Z-scores across all seven spermatid stages; Hi-C compartment scores (A compartment, positive; B compartment, negative); CUT&Tag read coverage (reads per million, RPM) for histone H4 at Earliest ES and Early ES (black); CUT& Tag read coverage for histone H2B at Earliest ES and Early ES (blue); and CUT& Tag read coverage for H4 acetylation at Earliest ES and Early ES (black). Heat map coloring is the same as Fig. 3A. **(C)** Median GC-regression-corrected ATAC-seq Z-scores across spermatid stages for genomic regions in the A compartment (solid purple) and B compartment (solid gold), with expected values from a standard normal distribution shown as dashed lines. Error bars represent 95% confidence intervals estimated by bootstrap resampling. **(D)** Immunofluorescence of individual spermatids isolated from specific stages of the seminiferous tubule cycle: Roman numbers are representative of the seminiferous tubule cycle stages, and Arabic numerals indicate spermatid steps in the 12 stages. Germ cells were co-stained for PRM1 (magenta, via V5 tag) and H4K20me3 (yellow), with Lectin (green). Scale bar: 2 µm.

When comparing genome-wide accessibility dynamics across compartments, we observe a clear difference in remodeling behavior at a low-resolution scale: A-compartment regions progressively lose relative accessibility as cells transition from late round to elongating spermatids, whereas B-compartment regions retain higher accessibility well into later elongating spermatid (Figure 6B). This compartment-specific pattern is observed across multiple chromosomes (Supplementary Figure 6C). Consistent with the representative chromosomal snapshots, the median accessibility levels for A-compartment loci drops quickly between the earliest and early elongating spermatid stages, whereas B-compartment regions retain higher accessibility well into late spermatids (Figure 6C). Notably, these distinct patterns observed are different from the expected background distributions, where both A and B compartments have median accessibility of around zero across all stages often with their confidence intervals overlapping (see **Methods**). To validate that the A compartment regions are likely to remodel first in vivo, we co-stained PRM1 with the heterochromatin mark H4K20me3 and monitored PRM1 localization across successive stages of elongating spermatids using structured illumination microscopy (SIM) (Figure 6D). In stage VIII–IX spermatids, PRM1 appears as discrete puncta localized to euchromatic regions and is clearly excluded from the H4K20me3-rich chromocenter. By stage IX-X, PRM1 spreads more broadly throughout the nucleus while still avoiding the heterochromatic center, and by stages XI–XII, PRM1 uniformly fills the elongating nucleus yet remains depleted from the chromocenter. Taken together, the in vivo and molecular data indicate that the timing of early versus late chromatin remodeling is strongly aligned with - and likely constrained by - the underlying 3D genome architecture established in round spermatids.

Consistent with these accessibility dynamics across compartments, A-compartment regions that lose chromatin accessibility during the transition from late rounds/earliest ES to Early ES also show parallel losses in relative histone H2B and H4 occupancy (Figure 6B). Tracking histone H4ac across compartments and stages reveals similar patterns. At late rounds/earliest ES (stages 7-8 spermatids), kernel density plots of H4ac suggest that H4ac levels are higher in A than in B compartments (Supplementary Figure 7A). However, this enrichment in the A compartment appears to be driven by the greater number of H4ac peaks in A compared with B (Supplementary Figure 7A). Furthermore, we find that the GC-normalized ATAC-seq distributions for H4ac-enriched loci in both compartments are remarkably similar and overlapping (histograms and box plots); suggesting that although H4ac sites are equally accessible in both compartments, they are selectively lost from the A compartment as cells enter Early ES. Consequently, by Early ES, the remaining H4ac signal is disproportionately retained in B-compartment regions (density plot; histogram; Supplementary Figure 7A). This apparent “shift” of H4ac toward B compartments is likely due to compositional consequences of differential histone loss and early closure of the A compartment. Consistent with this observation, in Early ES, the H4ac peaks that persist show notable differences in accessibility: those that persist in B compartments remain significantly more accessible than those few retained in A, indicating that once A-compartment chromatin begins its programmed closure, the presence of residual H4ac is insufficient to maintain accessibility.

### A proportion of H3K27me3-modified nucleosomes are stably maintained during spermiogenesis

A small proportion of nucleosomes is retained in mature sperm in a species-specific manner, with approximately 1-5% in mouse and 10-15% in human^14–16^. The large proportion of nucleosomes retained in mature sperm bear histone H3 lysine 27 trimethylation (H3K27me3), and include developmentally important loci, i.e., the Hox clusters, in both species^14–16,21^. Whether these retained nucleosomes reflect stable inheritance throughout the histone-to-protamine exchange process or are re-established at the end of spermiogenesis is unknown. To distinguish between these possibilities, we profiled H3K27me3-marked nucleosomes by CUT&Tag across Early RS, Late RS, Earliest ES, Early ES, and Int ES. Consistent with observations in mature sperm, we detected robust signal and enriched peaks at developmental promoters, including the *Hoxd* cluster; even when nucleosomes are destabilized (Supplementary Figure 7B; Supplementary Table 3). In contrast to H4-acetylated regions, which shift from A-to-B compartment in Early ES (Supplementary Figure 7A), H3K27me3-marked regions were maintained in the A compartment between spermatid transitions (Supplementary Figure 7C) and retained similar chromatin accessibility dynamics in both spermatid stages, unlike H4ac (Supplementary Figure 6A). To quantify the fraction of H3K27me3-marked loci stably maintained across spermiogenesis, we determined the proportion of peaks defined in Early RS that persisted through the ES stages. Approximately 60% of H3K27me3 peaks were retained (Supplementary Figure 7D), consistent with the possibility that these loci contribute to the retained nucleosome pool present in mature sperm. The removal of some H3K27me3-marked loci appears to be functionally important because mouse embryos generated by round spermatid injection (ROSI), which bypass the completion of histone-to-protamine exchange, exhibit ectopic retention of H3K4 and H3K27 methylation and show widespread misregulation of developmental genes, contributing to the low developmental competence of ROSI-derived embryos^52^. Complementary evidence from Xenopus has demonstrated that active demethylation of H3K4 and H3K27 in maturing spermatids is required for normal embryo viability^20^. Together, these findings suggest that H3K27me3 alone is insufficient to predict nucleosome inheritance in mature sperm; rather, the precise remodeling of both H3K4 and H3K27 methylation states during spermiogenesis is essential for establishing the epigenetic landscape required for embryonic development.

## Discussion

Protamines are the primary chromatin packaging proteins in sperm, yet, unlike histones, their deposition onto the paternal genome has been presumed to occur through nonspecific electrostatic interactions.^1^ Here we revisit this assumption and ask whether protamine-mediated genome packaging in sperm is random or programmed. Using endogenously tagged *Prm1* and *Prm2* mouse models, we demonstrate that the histone-to-protamine transition proceeds through a direct histone-to-PRM1 exchange, whereas TNPs precede only PRM2 incorporation. These findings demonstrate that PRM1 and PRM2 incorporation are temporally and, likely, mechanistically uncoupled, which revises the earlier model for Histone-to-Protamine exchange. To define the underlying molecular logic underlying the genome-wide chromatin remodeling, we mapped chromatin accessibility at high temporal resolution across spermatid maturation, and show that chromatin compaction is spatiotemporally asynchronous, proceeds through both transcription-dependent and transcription-independent mechanisms, and is facilitated but not directed by H4 hyperacetylation. Most strikingly, the order of compaction is strongly associated with the pre-existing 3D compartment architecture of the round spermatid genome. Together, these findings further underscore a programmatic nature and a compartment-correlated blueprint for establishing the paternal epigenome. (Figure 7)

**Figure 7:**
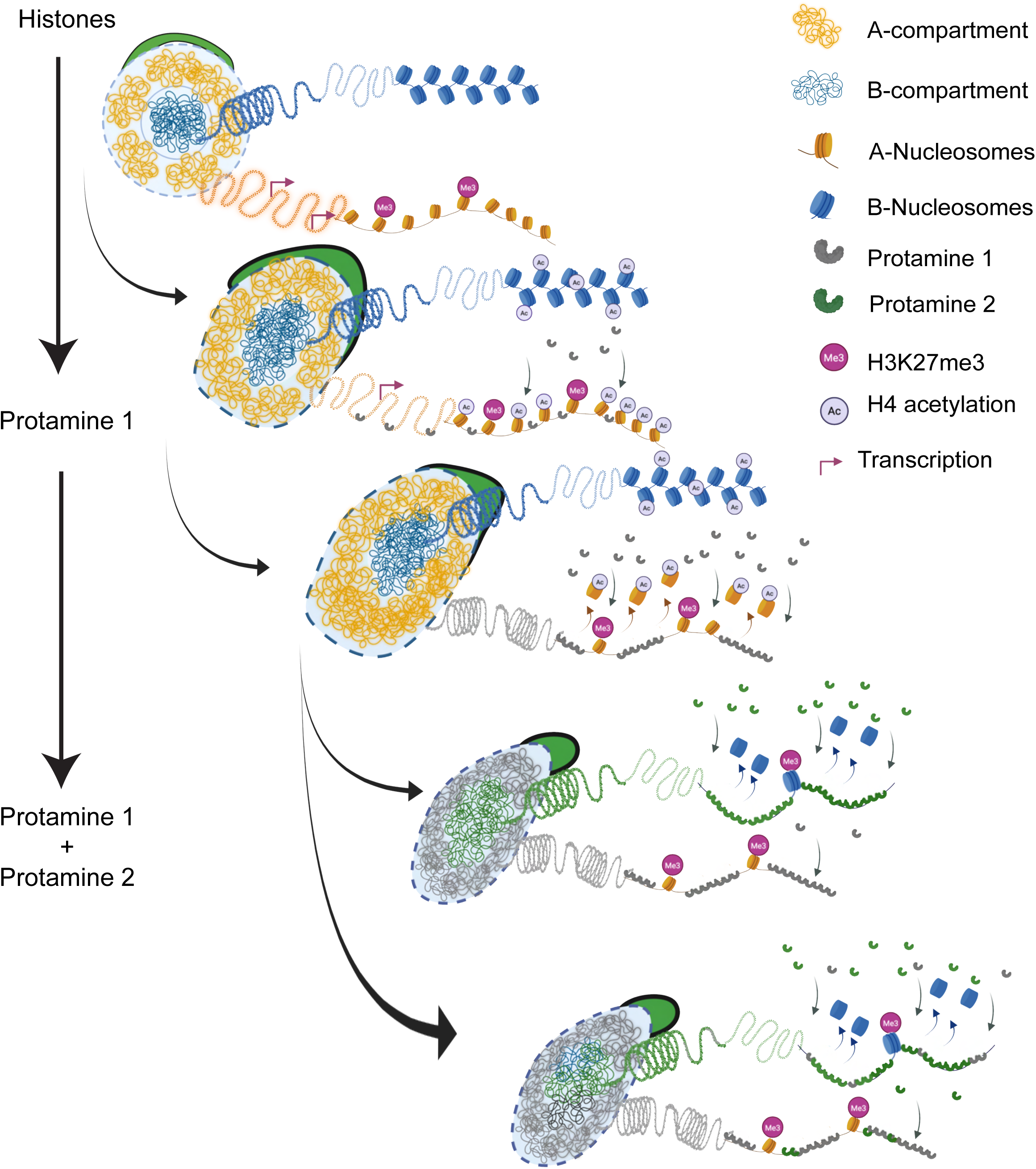
Model for compartment-encoded, programmatic histone-to-protamine exchange. This model illustrates a staged, compartment-resolved progression of chromatin remodeling during spermiogenesis. In late RS H4 hyperacetylation spreads globally, both in early and late remodeling loci, yet histone eviction and compaction initiate in the A-compartment chromatin, while B-compartment chromatin regions remain accessible until Late ES. Notably, A-compartment compaction coincides with PRM1 incorporation, whereas PRM2 appears late in spermatid development after TNPs, likely initiating a second, spatially distinct wave of compaction, potentially restricted to the B-compartment. The closure of the B compartment is predicted to be mediated either entirely by PRM2 or a combination of PRM1 and PRM2. Together, these findings define a programmed remodeling cascade in which the pre-existing three-dimensional nuclear architecture of the round spermatid nucleus dictates the order, timing, and spatial logic of histone eviction and protamine deposition.

The differential timing of PRM1 and PRM2 nuclear localization has been previously documented by antibody staining^7^ but has never been mechanistically interrogated or integrated into the canonical remodeling framework. Our endogenously tagged protamine models, combined with combinatorial staining, provide greater resolution of this revised histone-to-protamine exchange sequence. Genetic evidence supporting a direct histone-protamine1 exchange comes from *Tnp1/2* double-knockout mice, which showed normal PRM1 incorporation timing and levels, but also marked reductions in PRM2 incorporation, and, in some spermatids, a complete loss of PRM2^53^. This observed phenotypic difference in PRM1 and PRM2 regulation in these double TNP mutants can now be explained by the differential timing of PRM loading we observe and by the TNP-independent deposition of PRM1. This establishes PRM1 and PRM2 incorporation as mechanistically distinct events rather than steps in a single process. Further, this surprising, programmed uncoupling of PRM1 and PRM2 raises questions: do PRM1 and PRM2 occupy distinct genomic loci, and if so, what mechanisms direct each protamine to its target loci?

In early elongating spermatids, we find that PRM1 is recruited to H4-hyperacetylated chromatin. This early incorporation of PRM1 is likely mediated by BRDT, as deletions of the first bromodomain (BD1) have been shown to prevent the incorporation of PRM1^54–56^. Specifically, it has been proposed that nucleosomes bearing H4K5/K8 acetylation are recognized by BRDT, removed, and targeted for degradation by the PA200-containing proteasome^56^. Whether the BD1 domain of BRDT is similarly required for PRM2 incorporation remains to be determined; however, other histone modifications, such as H2BK12 crotonylation, have been implicated in TNP and PRM2 deposition in elongating spermatids^57^.

A key unresolved question is what regulates the stage-specific remodeling patterns we observe during spermiogenesis, since H4 hyperacetylation is broadly distributed across the genome in early and late remodeling regions as soon as it is established. Elegant work by Saadi Khochbin’s lab may provide insights into this sequence of remodeling events: H4K5/K8 butyrylation frequently co-occurs with H4 acetylation, and both exist independently at a subset of loc^56^. Early biochemical experiments show that the combination of H4ac and H4 butyrylation inhibits BRDT binding^56^. This suggests a combinatorial code in which acetylation, butyrylation, and crotonylation likely function as molecular timers that determine where and when structural exchange occurs, which inturn likely bias PRM1 vs. PRM2 incorporation.

By leveraging ATAC-seq insert size distributions, we provide insights into the underlying chromatin states during the histone-to-protamine transition. In early and Int-ES, we observed a marked loss of the canonical ∼147-bp nucleosome footprint, coincident with H4 hyperacetylation and the onset of PRM1 incorporation. Concurrently, the insert size distribution shifts towards a prominent ∼70-90 bp sub-nucleosome species, which is substantially wider than the TF-protected fragments or nucleosome-free region (NFR) in round spermatids. This sub-nucleosomal intermediate likely reflects destabilized nucleosomal states ^43^ or disassembly intermediates, such as hexasomes or tetrasomes, generated by the loss of one or both H2A-H2B dimer^58–60^. However, we also cannot exclude the possibility that some fraction of this signal reflects protamine-bound DNA fragments. Distinguishing between these possibilities will require orthogonal approaches, such as gentle MNase-based footprinting combined with nucleosome mass spectrometry ^61^or direct protamine mapping using single-molecule sequencingapproaches^62^. Both are extremely challenging due to cell number requirements ^61^ or technical reasons, due to the way protamine coats DNA^36^. However, the remodeling intermediate interpretation is nonetheless supported by evidence from other systems. In *Drosophila*, H2A-H2B removal precedes H3-H4 loss, generating a temporal series of disassembly intermediates during the transition to sperm nuclear binding proteins (SNBP)^59^. In *pal* mutants, H2A-H2B are removed normally, but H3-H4 tetramers are retained, yet this doesn’t impair SNBP loading, demonstrating that H3-H4 tetramers can coexist with SNBP-packaged chromatin^59^. A similar finding is observed in *Xenopus*, where H2A-H2B dimers are lost during spermatid maturation, but H3-H4 tetramers are selectively retained^60^. Whether these particles represent bona fide hexasome or tetrasome intermediates, protamine-bound DNA of similar fragment size, or destabilized full nucleosomes with impaired DNA wrapping - as proposed for remodeler-engaged nucleosomes in other systems - remains to be determined by systematic proteomic and single-molecule analyses.

A striking feature of early PRM1 incorporation is its punctate, euchromatin-restricted nuclear pattern, raising questions about how protamines first nucleate into these genomic regions. Our cross-correlations between ATAC-seq and scRNA-seq expression data across spermatid stages allow us to propose several potential models. In round spermatids, chromatin accessibility is tightly coupled to transcriptional activity, as expected. However, in Early ES, the most highly transcribed regions undergo a precipitous loss of relative accessibility - a transition that coincides precisely with the onset of punctate PRM1 incorporation. This could reflect a passive consequence of global chromatin reorganization, in which the rest of the genome becomes more accessible as elongation proceeds. However, we favor a more direct interpretation: that PRM1 is preferentially loaded into highly transcribed regions by virtue of their elevated accessibility because of nucleosome turnover or, more intriguingly, through transcription-coupled DNA strand breaks that create entry points for protamine intercalation. The latter is a more exciting possibility given that it has long been observed that post-meiotic round spermatids are known to have pervasive transcription^63,64^, yet its functional significance is unknown. Although prior assumptions suggest that this pervasive transcription is a ‘leaky’ byproduct of chromatin remodeling during spermiogenesis; we propose that this transcriptional activity is not incidental but is coupled to the initiation of histone-to-protamine exchange to establish a transcription-coupled chromatin remodeling pathway. Importantly, this mechanism accounts only for the remodeling of active genomic domains. The rapid and global decompaction of the B-compartment in Int- and late-ES suggests a parallel, distinct regulatory mechanism governing decompaction and re-compaction of heterochromatin. We speculate that this transition is driven by the programmed loss of heterochromatin-specific architectural factors, which, upon their removal, render these previously sequestered regions accessible to the nascent PRM 2 machinery. Thus, the genome-wide transition to a protamine-packaged state is a multi-step process: one governed by transcription-coupled remodeling in the A-compartment, and a second, independent pathway - potentially driven by the dissociation of repressive complexes - that triggers de-compaction across the B-compartment.

Whether protamines possess intrinsic DNA-sequence preferences or are guided to nucleation sites by dedicated chaperones remains unresolved. Protamines, like other basic proteins or zinc fingers, may have a modular ‘recognition code^65,66,67^. Indeed, simulation studies using arginine-rich short cationic peptides with varying numbers and spacing of consecutive arginine residues bias interactions toward GC-rich major-groove or AT-rich minor-groove in the double helix.^68^ Furthermore, in the real protein context in vivo, the protamine protein binding preference is also likely modulated by post-translational modifications that we and others have identified^69–71^. Once these proteins bind DNA, they appear to bind cooperatively^70,72,73^ – beginning with a nucleation event that spreads outward - a behavior supported by our earlier in vitro biophysical assessments on DNA curtains^70^ and by our imaging data here. This cooperative spreading is consistent with a bind-and-bend mechanism, in which initial protamine binding induces a local DNA conformational change that increases the probability of subsequent protamine binding and loop expansion.^72^ Consistent with this, we observe dense clusters of PRM1 in early spermatids, which become more diffuse in later stages. However, stably retained nucleosomes, particularly those marked by H3K27me3, could act as barriers to protamine domains, a finding consistent with previous models.^36^

Although locus-specific genomic/biochemical cues clearly shape where and when protamines load, our data reveal a broader organizational principle that correlates with and likely reflects the re-established A/B compartment structure of the round spermatid genome. This compartment structure, which is reinstated following its dissolution during meiosis, serves as a powerful predictive blueprint for how chromatin is remodeled: i.e., A-compartment regions compacting earlier than B-compartment regions, creating a spatially coherent, genome-wide progression that can’t be driven by local histone modification alone. This re-established chromatin compartment identity in rounds is not a transient feature of spermiogenesis; it is maintained/strengthened in mature sperm and directly passed on to the paternal pronucleus of the one-cell zygote.^74,75–77^ These observations suggest that sperm can deliver more than DNA methylation and a small fraction of retained histones, but rather provide compartment-level information to the zygotes that is encoded by PRM1 vs PRM2. In line with this observation, it has been shown in mouse embryos that the paternal pronucleus has distinct chromatin compartments, whereas maternal pronuclei do not.^76^ Furthermore, the emerging compartment landscape, starting with the two-cell embryo, appears to follow the compartmental patterns observed in mature sperm, suggesting the exciting possibility of a paternal epigenome-guided chromatin organization in the embryo.^75^ Furthermore, it has also been shown that A and B compartments precede, guide zygotic transcription, and instruct replication timing in early embryos.^78^ Together, these observations suggest that higher-order chromatin organization established in spermiogenesis could have implications in early embryo development.

## Acknowledgments

We thank all members of the Hammoud lab for their discussion and feedback on the manuscript. We thank Christa Ventresca, Lovelyn Epelle, Catherine Tower, Dominic Bazzano, Noah Helton, Ray Trievel, Nicholas Bockhaus, University of Michigan Flow Core, University of Utah Genomics Core (Brian Dalley; HCI’s Cancer Center Support Grant), and University of Michigan Transgenic Facility. This research was supported by National Institute of Health (NIH) grants GM148028 (S.S.H.), 1DP2HD091949-01 (S.S.H.), R01HD104680 01 (S.S.H.), R01HD113274 (S.S.H.), GM148028, T32 Program 5T32HD079342-10 (D.B.), CTRB training grants T32-HD079342 (S.S.), University of Michigan Pioneer fellowship (R.A.) and Open Philanthropy Grant 2019-199327 (5384) (S.S.H.).

## AUTHOR CONTRIBUTIONS

S.S.H and M.R. conceived and designed the experiments. Experiments were performed by M.R., L.M., S.S., R.A., W.X. Data analysis was done by T.E.W, Z.A., S.K., T.P., M.R, M.K., A.V., P.O. Computational Advice was received from J.Z.L., A.P.B., L.F. Essential Experimental Reagents were obtained from B.L. and B.C. The manuscript was written by S.S.H. and M.R. with input from all authors.

## Methods

### Mice

All mouse (*Mus musculus)* experiments were carried out with approval from the University of Michigan Institutional Committee on Use and Care of Animals (PRO00006047, PRO00008135, PRO00006047, PRO00008135, PRO00010000, PRO00011691), in accordance with the guidelines established by the National Research Council Guide for the Care and Use of Laboratory Animals. Mice were kept at the University of Michigan animal facility in an environment with controlled light (12 h on/off), temperature (21-23 °C), and humidity (30–70%), with ad libitum access to water and food (Lab Diet no. 5008 for breeding mice, no. 5LOD for non-breeding animals).

PRM1^+/V5^ mice, PRM2^+/V5^ mice were generated on a mixed genetic background (BL6 / SJL hybrid) using CRISPR–Cas9-mediated genome editing. Transgenic mouse models were generated by the University of Michigan Transgenic Animal and Genome Editing Core Facility. All animal procedures were carried out in accordance with guidance from the Institutional Animal Care and Use Committee and approval from the University of Michigan Medical Center. All mice were backcrossed to the C57BL/6J background for at least 6 generations.

### Assessment of fertility of mutant mouse models

To assess fertility, epididymal sperm were collected in 2ml of Donner’s media (25 mM NaHCO_3_, 20 mg/mL Bovine Serum Albumin, 1 mM sodium pyruvate, 0.53% vol/vol sodium DL-lactate in Donner’s stock)^1^ for 30min at 37°C. Briefly, the collected sperm were used to assess sperm count, motility, and morphology. Sperm count and motility were measured using a Makler chamber. Measurements were taken in triplicate and >100 sperm counted for motility.

### Fertility assessment

Fertility assessment was done as previously described^2^. Briefly, male mice (8 weeks old) were individually housed for 3 days before mating. Each mail was then paired with three females for three independent replicates per genotype. Females were checked daily for copulatory plugs and once detected, were separated into individual cages. Fertility was assessed by recording the number of females successfully impregnated.

### Immunofluorescence (IF) and seminiferous tubule staging

Testes were collected and fixed in 4% PFA in 4°C 4°C overnight with rotation, washed in 1X PBS overnight, and stored in 70% ethanol prior to formalin-fixed paraffin embedding (FFPE). Embedded tissues were sectioned into five-micrometer-thick tissue sections and deparaffinized and permeabilized in 0.1% triton X-100 in PBS for 15min. Tissue were then subjected to antigen retrieval by boiling in 10 mM sodium citrate, pH 6.0, for 30 min, followed by 20 min of cooling at room temperature while submerged in the sodium citrate solution. Sections were blocked in 3% BSA in 1X PBS for one hour, followed by incubation in wash buffer (3% BSA, 500 mM glycine in 1X PBS) for two hours and a 10-minute block in Bloxall. Primary antibodies were applied overnight at 4°C. Sections were washed four times in PBST (0.1% BSA, 0.05% Tween-20 in PBS) prior to incubation with secondary antibodies (1:500; Life Technologies/Molecular Probes). PNA-lectin (1:500; GeneTex) and DAPI were added simultaneously with secondary antibodies to label the acrosome and total nuclear DNA, respectively.

For the detection of mouse primary antibodies against Hup1n, Hup2b, and Tnp2, immunofluorescence was performed using the M.O.M. Mouse on Mouse Kit (Vector Laboratories) according to the manufacturer’s instructions, with the following modifications: primary antibody incubation was extended to overnight, and sections were washed four times in PBST prior to secondary antibody addition.

Seminiferous tubule stages were assigned to cross-sections based on PNA-lectin staining patterns of the spermatid acrosome and the complement of cell types present within each tubule, categorizing tubules into stages VIII, IX–XII, I–III, and IV–VI as previously described.^3^

### Antibodies

For IF staining, antibodies were used at the following dilutions, Hup1n: 1:100, Hup2b:1:100, mouse V5: 1:500, rabbit V5: 1:100, TNP1: 1:100, TNP2: 1:50, histone H3: 1:500, histone H4ac: 1:500.For western blot, PRM1 custom ab was used at 1:500 and histone H3 at 1:1000. For Cut&Tag, antibodies were used in the following dilutions: H4: 1:100, H2B: 1:50, H4ac:100, H3K27me3:1:100.

**Catalog no of antibodies used:**

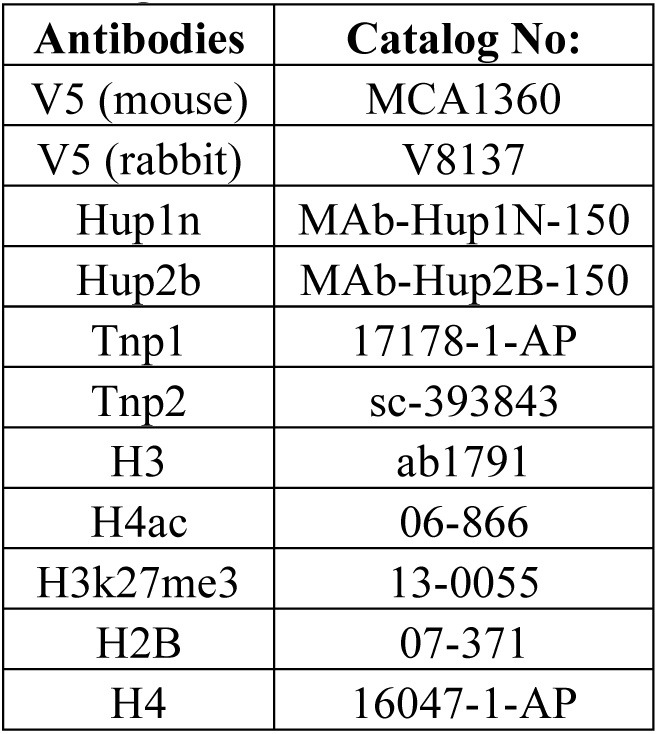

### Synchronization of spermatogenesis in juvenile mice

Spermatogenesis was synchronized in male pups by feeding them a retinoic acid inhibitor WIN 18446, starting at 2 days post-partum for 7 days followed by an injection of retinoic acid dissolved in DMSO, as previously described ^4^ with slight modifications. Retinoic acid dosage was reduced from 200ug to 100ug to reduce retinoic acid-mediated toxicity. Testes were collected beginning with the second wave of spermiogenesis from day 31 onwards. For each animal, 1.5 testis was processed for experimental analysis while the remaining ½ was reserved for stage verification. Synchronization was confirmed by PNA-lectin staining of acrosome morphology on cross-sections of the reserved testis, ensuring that all samples collected and analyzed corresponded to the intended spermatogenic stage.

### Salt fractionation of chromatin

Testes were synchronized as described above and collected from the second wave of spermiogenesis, after retinoic acid injection. Snap-frozen testes were homogenized in a Dounce homogenizer, washed with PBS, and incubated with 0.1% CTAB on ice for 5 minutes to remove flagella. Cells were washed twice in 50 mM Tris-HCl (pH 8.0) and lysed on ice for 15 minutes, then centrifuged at 800g for 15 minutes. The supernatant was retained as the cytoplasmic fraction. Pelleted nuclei were digested with 1 U MNase at 37°C for 30 minutes, quenched with EGTA, and centrifuged at 400g for 10 minutes; the supernatant was collected as the MNase fraction^5^. Remaining chromatin was subjected to sequential washes with 0.5 M, 1 M, and 2 M NaCl for 30 minutes each^5^, with supernatants collected after centrifugation at 400g for 10 minutes following each wash. Proteins in each fraction were precipitated by addition of 20% TCA and overnight incubation at −20°C with rotation, pelleted at 12,000g for 10 minutes at 4°C, and resuspended in water prior to immunoblotting.

### Isolating spermatids from specific stages for ATAC-seq

#### Single cell dissociation

Testes from adult mice were collected from adult mice as previously described^6,7^, and the tunica albuginea was removed. S Seminiferous tubules were transferred to 10 ml of Digest Buffer 1 (Advanced DMEM: F12 supplemented with 2 mg/ml Collagenase IA (Sigma) and 0.2 mg/ml DNase I (Worthington Biochemical Corp)), inverted 5–10 times, and incubated horizontally at 35°C with shaking at 215 rpm for 5 minutes. Tubules were allowed to settle for 1 minute at room temperature and the supernatant was discarded. Tubules were then resuspended in 10 ml of Digest Buffer 2 (0.5 mg/ml Trypsin and 0.4 mg/ml DNase I in Advanced DMEM: F12), inverted 5–10 times, and incubated at 35°C with shaking at 215 rpm for 5 minutes. Tubules were further dissociated by pipetting 3–5 times with a 1 ml pipette tip, and trypsin activity was quenched by addition of 3 ml fetal bovine serum (FBS; Sigma). The cell suspension was filtered through 70 micrometer strainer and centrifuged at 300g for 5min at 4 °C. Cells were then resuspended in 6ml of MACS buffer containing 0.5% BSA (Miltenyi Biotec).

#### Fluorescent activated cell sorting

Dissociated cells in MACS buffer containing 0.5% BSA were quantified using a hemocytometer. 0.06μL (0.6μg) of Hoechst 33342 dye was added per 1million cells alongside 4μL of Syto 16 dye. Cells were shaken in the incubator at 35 °C/215rpm for 20min. An equal amount of Hoechst 33342 and Syto16 dyes was added again, and cell suspensions were shaken in the incubator at 35 °C/215rpm for another 20min. Cells were then centrifuged at 300g for 5min at 4 °C and resuspended in 6-10ml of MACS buffer. 10ul of Draq7 dye was added, and cells were filtered through 70 micrometer filters and used for FACS. Cells were gated as previously described in Gill et al. 2022^8^ to sort for round spermatids and elongating spermatids.

### ATAC-seq experiments and sequencing

ATAC-seq was performed as described in Corces et al. 2017^9^ with minor modifications. Briefly, FACS-sorted spermatids were centrifuged at 500g for RS and 2500g for ES for 10 minutes at 4 °C. 220,000 cells were resuspended in DnaseI buffer (500ul 1X HBSS, 2.5ul of 1M MgCl2, 0.5ul of 1M CaCl2) with 2.5ul of 20mg/ml DNaseI. Cells were treated at 37 °C for 20min. Cells were washed with 1X PBS twice during the second wash, 10% Drosophila spike-in cells were added prior to centrifugation. Cells were resuspended in ATAC resuspension buffer (10 mM Tris-HCl pH 7.5, 10 mM NaCl, 3 mM MgCl₂) supplemented with 0.1% IGEPAL, 0.1% Tween-20, and 0.01% digitonin, and monitored for lysis by trypan blue exclusion. C Lysis was quenched by addition of 900 µl ATAC resuspension buffer containing 0.1% Tween-20, and nuclei were pelleted at 2,500g for 10 minutes at 4°C. Transposition reaction was performed by resuspending nuclei in transposition mix (50 µl 2X TD buffer, 5 µl transposase, 33 µl PBS, 1 µl 1% digitonin, 1 µl 10% Tween-20, 10 µl H₂O) and incubating at 37°C for 30 minutes with 1,000 rpm mixing. Reactions were cleaned using Qiagen MinElute Kit according to the manufacturer’s instructions. Libraries were amplified as previously described^9^ with 5 additional cycles in the final amplification step. Final libraries were purified with the Qiagen MinElute Kit, and further size-selected using 1.5X AMPure XP beads. Final libraries were sequenced as 150-bp paired-end reads on an Illumina NovaSeq X. Note: later-stage spermatids beyond stage 4 were very difficult to permeabilize and generate reliable samples without significantly perturbing chromatin. We therefore opted to stop at these stages. We know DTT treatment in sperm affects the DNA binding ability of certain modified protamine molecules.

### CUT&Tag experiment and sequencing

CUT&Tag protocol for spermatids was adopted from EpiCypher CUTANA^TM^ Direct-to-PCR CUT & Tag Protocol and Kaya Okur et al 2019^10^ with modifications. Briefly, FACS-sorted spermatids were centrifuged at 500g for RS and 2500g for ES for 10 minutes at 4 °C. 220,000 cells were resuspended in DnaseI buffer (500ul 1X HBSS, 2.5ul of 1M MgCl2, 0.5ul of 1M CaCl2) with 2.5ul of 20mg/ml DNase I. Cells were treated at 37 °C for 20min. Cells were then washed twice with 1X PBS. Round spermatids were lysed with NEBuffer + 0.1% triton 10min on ice. For elongating spermatids, cells were treated with 0.05% lysolecithin for 10min. Cells were then resuspended in NEBuffer + 0.1% Triton X-100 + 0.1% digitonin and monitored for lysis using trypan blue exclusion. 100ul nuclei were aliquoted into 10ul of activated conA beads in NEBuffer + 0.1% triton and incubated together for 10min at room temperature. Nuclei were incubated with primary antibodies overnight on a nutator. Nuclei were incubated with species-specific secondary antibodies at ®a 1:100 dilution for 1 hour at room temperature. All subsequent steps were done according to EpiCypher CUTANA^TM^ Direct-to-PCR CUT&Tag Protocol. All libraries were amplified using 14 PCR cycles. Libraries were cleaned with 1.3X AmpureXP beads and sequenced with 150 bp paired end reads on an Illumina NovaSeq X.

### Hi-C experiment

One million FACS-sorted round spermatids were centrifuged at 500g for 10min at 4°C. Cells were resuspended in 1ml PBS and 2% formaldehyde. All subsequent steps were according to the Arima Hi-C protocol (Arima HiC-Kit, User Guide Mammalian Cell Lines, Doc A160134 v01). Libraries were generated across 4 biological replicates and sequenced using 150bp paired end reads on an Illumina NovaSeq X.

## DATA ANALYSIS

ATAC-seq data generated across the seven key stages of spermiogenesis were processed in parallel using two different pipelines. Key differences between the two methods are outlined below and will be referred to as Pipeline 1 and Pipeline 2.

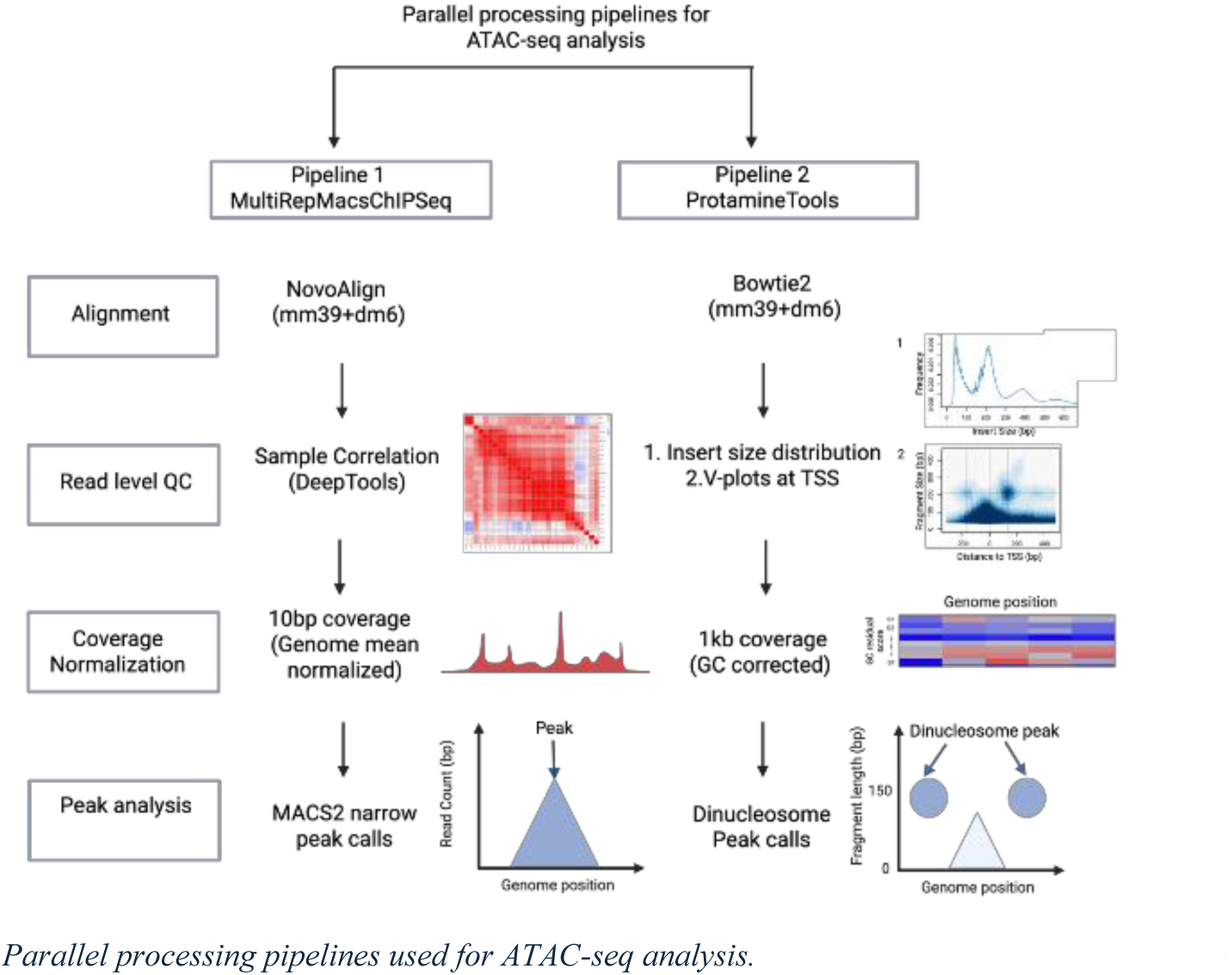

### ATAC-seq analysis – Pipeline1: MultiRepMacsChIPSeq

#### Alignment and Normalization

ATAC-seq reads were aligned using Novoalign (version 4.04.01, http://novocraft.com) to a combined mouse (UCSC version mm39) and *Drosphila* (UCSC version BDGP6) reference genome using default-tuned parameters for NOVASEQ and including built-in adapter trimming with sequence CTGTCTCTTATACACATCT. Alignments were tagged with mate scores using Samtools fixmate (version 1.21, https://www.htslib.org/doc/1.21/samtools-fixmate.html). Alignments were split into separate genome-specific files using Picard ReorderSam (version 2.23.3, https://broadinstitute.github.io/picard/). Alignment files from samples and replicates were put through the MultiRepMacsChIPSeq pipeline (https://huntsmancancerinstitute.github.io/MultiRepMacsChIPSeq/, release 20) for fragment-based peak calling. Properly-paired alignments were filtered based on insertion length (40-300 bp), exclusion of mitochondrial and contig chromosomes, removal of optical duplicate alignments (pixel distance of 250), and partial PCR de-duplication via random subsampling to a final total duplication rate of 5%. Genome-wide fragment coverage was generated (in 10 bp bins), normalized to a depth of 1 million fragments. Coverage was scaled to the median sequencing depth of the given samples, statistical enrichment (q-value) genome-wide tracks generated using MACS2^11^ and the empirical genomic mean coverage, and peaks called using MACS2^11^ using a minimum peak size of 200 bp, gap of 50 bp, and minimum threshold of 0.01 (1% FDR).

Delta coverage tracks were generated by taking bedgraph file format of the coverage tracks using MACS2^11^ (version 2.2.7.1) bdgcmp function to substract the genome mean of the sample, which was calculated using the custom script generate_mean_bedGraph.pl (https://github.com/tjparnell/HCI-Scripts/blob/master/ChIPSeq/generate_global_mean_bedGraph.pl). The resulting bedgraph files were visualized by converting to bigwig files. Correlation heatmap between aligned bam files were generated using the function multibamsummary of deeptools^12^ (version 3.5.4) and python (version 3.11.5) with a bin size of 10kb.

#### Gene ontology analysis

MACS 2 peak calls per stage were merged using bedtools2^13^ (v2.31.1) ‘merge’ and intersected with transcriptional start sites of mm39 reference genome using bedtools2^13^ (v2.31.1) ‘intersect’. The resulting unique gene names were used to perform gene ontology analysis using David Bioinformatics^14^ (https://davidbioinformatics.nih.gov/workspace.html).

#### ATAC-seq analysis – Pipeline2: Protamine-tools

ATAC-seq data from spermatids presents distinct challenges relative to the application of the method to other cell types because chromatin states are altered on a genome-wide scale during spermiogenesis. Regions with increased insert counts can represent both typical positioned nucleosomes with localized hyper-accessible nucleosome-free regions, e.g., those associated with changing transcriptional activity, as well as extended highly accessible regions during protamine deposition. We created new tools for integrating this unique data class with other data types in the ‘protamine-tools’ pipeline (https://github.com/HammoudWilson/protamine-tools). The workflow is divided between stage 1 data analysis pipelines and stage 2 interactive plotting apps created using R Shiny. Detailed instructions are provided, although much of this purpose-specific codebase will only apply to similar spermatid data sets. The codebase includes normalization approaches that were initially explored but were not used in any presented figures; these are discussed more briefly below.

##### Read alignment to a mouse-fly composite genome

We first created a composite mouse-fly FASTA file with chromosomes from the mm39 and dm6 reference assemblies using protamine-tools actions ‘genome download’ and ‘genome compositè. Chromosome names were suffixed with the genome as ‘chr1_mm39’, etc. Read pairs were prepared for alignment by ‘atac align’ by trimming them to 150 bases per read followed by quality filtering and merging of overlapping read pairs using ‘fastp’ v0.24^15^ with settings ‘--dont_eval_duplication --length_required 35 –correction --merge --overlap_len_require 20 --overlap_diff_limit 10 --overlap_diff_percent_limit 20 --trim_poly_g --adapter_sequence CTGTCTCTTATACACATCT’. Prepared reads were aligned to the composite genome using ‘bowtie2’ v2.5 with setting ‘-x -U --end-to-end’ for merged single reads and ‘--end-to-end --maxins 650 --no-mixed --no-discordant’ for read pairs from longer inserts and alignments merged to a single BAM file. Merged reads and concordant read pairs were deduplicated such that no two DNA inserts shared the same alignment endpoints. Reads were filtered to require a minimum mapping quality (MAPQ) of 30 to ensure high confidence placements.

##### Binned data analysis

Due to the rapidly changing accessibility landscape of large genomic regions during spermiogenesis, we first assessed ATAC-seq insert yield at low resolution by binning the composite genome into 1kb bins using the pipeline action ‘genome bin’. The approach sought to assess regional Tn5 accessibility. Throughput, bins in the mm39 and dm6 GENCODE blacklists^16^ were masked in files as “not included” in statistics and plots. Various genome and sample level scores were collected for each bin for correlation and plotting as described below.

##### Raw bin counts and insert size distributions

Filtered and deduplicated ATAC-seq inserts were counted in the bin that contained their leftmost aligned ends by pipeline action ‘atac collatè. Insert counts were normalized to reads per kb per million reads (RPKM) for inter-sample comparison. RPKM is synonymous with count per million reads (CPM) at 1kb bins, but bins were often aggregated further below. Due to the relative sizes of bins and inserts, especially aggregated bins, most inserts were fully contained in one bin or region. Some inserts crossed into adjacent bins, which was tracked during next steps. Insert size distributions were assembled per sample library and per genome, so that insert sizes could be compared as a function of both spermatid stage and origin from the mouse vs. Drosophila spike-in DNA.

##### Mappability and other insert size-dependent scores

An unusual feature of our ATAC-seq data is that the sequenced insert sizes vary considerably across different spermatid stages, leading to potential latent insert size-dependent biases during sample comparison. We assessed the mappability of the composite genome as a function of insert size from 35 to 650 bp, the range used for ATAC-seq analysis, using action ‘genome mappability’ in a manner that considered insert size, since small inserts are inherently less mappable in repetitive sequences. Insert sizes were stratified into twenty groups from beginning at 35, 40, 45, 50, 55, 60, 70, 80, 90, 100, 120, 140, 160, 180, 200, 220, 240, 260, 280, 300 bp. The ‘genmap map’ utility, v1.3^17^, was used to calculate the unique mappability of inserts at the smallest insert size in each group with settings ‘-K <INSERT_SIZE=-E <N_ERRORS> -bg’, where n_errors was 1 for insert_size <= 40, 2 for insert_size <= 100, 3 for insert_size <= 200, and 4 for insert_size >= 200, to allow for increased numbers of sequencing errors in longer inserts. Bin mappability was calculated on a per sample basis as an average of the mappability of the different insert size levels weighted by the sample insert size distribution. A bin’s raw insert count was divided by the dynamically calculated mappability to yield adjusted insert counts. Bin counts were rescaled to the same summed count after mappability correction to avoid inflating observed counts.

We explored using insert size profiles as a primary bin score. Specifically, we used samples from early RS and intermediate ES stages to establish emission probabilities for nucleosome-bound and transitional and/or protamine-bound states, respectively. We calculated the relative log likelihood for each bin for the early and late-stage emission profiles and normalized it to the bin count. The resulting normalized relative log likelihoods (NRLL) were visualized in R Shiny but not depicted in figures as the information proved redundant and less stable than other scores.

##### GC bias normalization and GC-related bin properties

Tn5 libraries have a notable bias in mapped read recovery based on local base content. The extent of this GC bias differed significantly between ATAC-seq samples collected at different spermatid stages, in part due to their different insert size distributions since smaller inserts tend to be more GC rich. For low-resolution analysis this technical effect becomes a substantial determinant of bin coverage, although some component of true biological differences between samples may correlate with GC content.

The GC base content of bins was assessed for DNA inserts that could productively map to each bin for each insert size group and a net effective GC value calculated for each sample-bin as for mappability above. We correlated mappability-adjusted bin insert counts to this bin GC content and performed regression analysis using the negative binomial distribution using R expression ‘MASS::glm.nb(formula, control = glm.control(maxit = 100))’, where formula was ‘readsPerAllele ∼ fractionGC + fractionGC2’ and ‘fractionGC2’ was the square of the bin GC fraction to achieve a quadratic fit of the trend toward higher coverage at higher GC content and the degree of count over-dispersion. We calculated the “GC residual Z-score” (GCRZ) as the deviation from the trend line in Z units by projecting the negative binomial quantiles onto a normal distribution. Thus, samples with a high GCRZ were overrepresented relative to peer bins with a similar insert size-adjusted GC content and runs of bins with high GCRZ scores were judged to be enriched independently of bin GC content. For regression we only considered bins with GC fraction between 0.25 and 0.6 to prevent outlier effects from rare bins with extreme values.

Two additional factors were considered as contributors to the strong GC bias in spermatid ATAC-seq. CpG dinucleotides are more frequent in higher GC content bins. Indeed, the relative insert yield in bins with high CpG content changed substantially as spermiogenesis progressed and signal shifted away from gene promoters toward the rest of the genome. However, this effect was not a substantial contributor to the changing GC bias across stages since bins with high CpG content are a small fraction of the genome and those bins went against the trend of higher GC preference in later spermatid stages. Tn5 also has a well-described base preference, although not a strict requirement, that leads to non-random base utilization at insert ends^18^. Its preferred motif is GC-rich, an effect which becomes increasingly important as insert size decreases. Pipeline action ‘atac sites’ used observed inserts across all samples to establish a discontinuous 9-based Tn5 preference motif at the bases with the strongest Tn5 preference, 5’-**|*-***-***, where* is a preferred base and | is the cleaved bond. Bins were assigned weights based on their relative enrichment for preferred sites, and the GC bias fit was repeated with weight-adjusted insert counts. GC bias was lower after independently accounting for Tn5 site preferences, but the net results did not change, presumably because the unweighted analysis already accounted for this effect.

##### Transcription level of bins and annotated transcription start sites

We used mouse round spermatid PRO-seq data^19^, obtained as BigWig files from GEO series GSE228452, as our baseline to understand the transcription levels of genes and bins. Data representing the 3’ terminus of PRO-seq reads were lifted over from the mm10 to mm39 reference assemblies, aggregated over both strands, counted per bin, and normalized to yield log10 RPKM by pipeline action ‘nascent bin’. Bins with PRO-seq signal below a log10 RPKM of -3 were having no evidence of transcription. Separately, we examined the 5’ ends of genes in the GENCODE M36 mouse genome annotation, i.e., putative transcription start sites (TSS), for evidence of transcriptional activity and positioned nucleosomes to use as references points for chromatin changes during spermiogenesis. TSSs were scored per-sample as “active” by pipeline action ‘nascent tss’ when the 1kb downstream of the 1^st^ base of the gene had a PRO-seq RPKM of at least 0.5 and there was at least a 10-fold increase in RPKM when compared to the 1kb upstream of that base. Genes whose 1^st^ 1kb had RPKM < 0.05 were considered inactive for TSS assessments. For active TSS, we attempted to find evidence for positioned a positioned nucleosome just downstream of the TSS using pipeline action ‘atac tss’ when there were at least 25 mononucleosome-sized ATAC-seq inserts (150 to 275 bp) covering the region within 250 bp of the gene start. The position of the inferred position nucleosome was taken as the median of the insert midpoints. Metadata plots over all active or inactive TSS were registered based on either the position of the TSS or the inferred nucleosome center.

##### CUT&Tag score integration

BigWig files from H2B, H4, H3K27Me3 and H4Ac CUT&Tag data were handled in a manner like PRO-seq data by summing insert counts over both strands per bin in pipeline action ‘cuttag scorè. Here, for some histone targets many or most bins had an RPKM of zero.

##### Bin aggregation to lower resolution plots (horizontal aggregation)

Many plots and analyses compared wider genome regions at considerably lower resolution than 1kb bins. Horizontal bin aggregation was achieved by summing the counts or averaging the normalized scores, such as RPKM, of the 1kb bins that crossed each pixel in the final image, weighted by the fraction of the bin overlapping that pixel. Thus, each pixel faithfully reports the aggregated data it contained at the resolution of the image.

##### Sample aggregation into spermatid stages and stage types (vertical aggregation)

Working from prior knowledge of the biological properties of the spermatid samples and visual examination of ATAC-seq data, we defined seven time-ordered stages of spermiogenesis: early round spermatids (RS; early_RS), intermediate RS (int_RS), late RS (late_RS), earliest elongating spermatids (ES; earliest_ES), early ES (early_ES), intermediate ES (int_ES), and late ES (late_ES). RS stages showed differences mainly in the location of typical chromatic ATAC-seq peaks, earliest_ES behaved as a transition stage, and the later ES stages showed large differences from typical ATAC-seq manifest as broadly increased accessibility and smaller insert sizes. Accordingly, we further defined two stage types, RS and ES, that grouped the stage types with matching round/elongating labels. To simplify plots, we performed a vertical aggregation of at least two samples per stage or stage type by summing ATAC-seq insert counts prior to making score calculations by the same methods as for individual samples. We finally defined a stage type delta as RS – ES to yield a single score per bin with high values when RS had more signal than ES.

##### Stage mean calculation

We devised a further way of assessing bin scores, especially ATAC-seq RPKM, that considered all samples together in a single bin or region to reveal which stages along the timed series were most enriched for higher scores. For this “stage mean” metric, we indexed the spermatid stages from 1 (early_RS) to 7 (late_ES) and took an average of the stage indices weighted by the average score values of each stage’s samples. Thus, low and high stage means indicate that early or late-stage samples, respectively, showed higher signal relative to the rest of the genome in that bin or region. Because mappability and GC biases are largely the same across samples in a bin or region (aside from insert size-dependent effects noted above), stage mean values are robust to local genomic effects and provide a reliable score for the timed changes occurring during spermiogenesis. Stage mean scores cannot distinguish U-shaped patterns; higher scores in both early and late stages will give a middle stage mean value. However, most bins showed either decreasing or increasing scores, or peaks centered on a specific stage, where stage mean is a faithful reporter.

##### Normalization for heat map plotting

Heatmap plots used a color scheme where red colors were “hot” and correlated with increased signal, or greater signal in round vs. elongating spermatids, and blue colors were “cold” and the opposite of red/hot. Some scores were inherently centered and scaled and required no further normalization, including GCRZ and Hi-C compartment scores. PRO-seq log10 RPKM represents a varying intensity of positive signals so only red/hot colors were used to reflect the level of local transcription up to a fully red color at a log10 RPKM of 10. For other scores, heatmap plots were normalized using Z-scores of normally distributed score values, e.g., bin GC fraction, or as quantiles of non-normal score distributions centered around the median. Importantly, most ATAC-seq, CUT&Tag and Hi-C scores are relative values across the genome.

##### Base-resolution positioned dinucleosome peak calling

Most spermatid stages also revealed changing local accessibility peaks associated with positioned nucleosomes that we sought to track. Many algorithms are challenged in finding these peaks in samples with much broader accessibility, where accumulations of mapped inserts were often called as peaks that did not represent the same biological mechanisms as accessibility peaks at promoters, enhancers, and other functional genomic elements. We developed an efficient base-level algorithm in Rust to locate “dinucleosome peaks” characterized by (i) two nucleosome-size spans frequently crossed by longer ATAC-seq inserts but with few endpoints mapping within them, and (ii) intervening and flanking nucleosome-free regions with a higher density of insert endpoints and coverage by smaller inserts. Such peaks are distinct from other generally hyper-accessible regions in showing a specific insert pattern that is evident on V plots and characteristic of genome elements that coerce nucleosomes to specific locations.

Peaks were called *ab initio* working only from ATAC-seq insert endpoints. The algorithm finds the most likely positions of positioned dinucleosomes in local regions, if there were any, and filters those positions against minimal criteria for dinucleosome peak calling. Working per chromosome, three base-level maps of inserts locations are accumulated over all samples of a specific spermatid stage. The first map counts the number of insert endpoints at every base. The second counts the centers of mononucleosome-size inserts from 147 to 320 bp. The third counts the centers of dinucleosome-size inserts from 320 to 485 bp, which are expected to fall in the nucleosome-free regions between two positioned nucleosomes. A series of counting masks represent different candidate gap lengths of the intervening nucleosome-free region from 51 to 351 bp with a step size of 4. Nucleosomes occupy 147 bp flanking each gap, and an additional 75 bp are included on each nucleosome flank as regions expected to have high endpoint counts.

For every third base along the chromosome, the insert maps and counting masks are used to generate a positive score where (i) insert endpoints within nucleosome-free bases are counted with a weight of 1, (ii) mononucleosome-size insert centers in nucleosome-bound bases are counted with a weight of 2 (the weight of two endpoints), and (ii) dinucleosome-size insert centers in nucleosome-free bases are counted with a weight of 1 (making them less important than mononucleosomes). In this way, many inserts will not be counted when the gap length is sub-optimal or nucleosomes are not positioned as endpoints and centers will not conform to the mask. The highest scoring combination of genome position and dinucleosome spacing establishes a local maximum that is called as the first peak. No other combination in the vicinity that would override the gap of the first, highest scoring combination is considered further. The portions of the chromosome flanking each peak are then analyzed recursively in the same way until no additional high-scoring peaks remain.

After calling peaks for a spermatid stage, adjacent peaks that overlap in their nucleosomes or flanking nucleosome-free regions are merged into a “nucleosome chain”, i.e., a span with three or more positioned nucleosomes, by taking an average of the nucleosome positions and gap lengths weighted by the peak scores. Chains grow iteratively until no more overlaps are found. Chains are then intersected between all samples to find “peak regions” as the widest coordinate span of all overlapping chains from different stages. RPKM and stage mean values are calculated for all samples for each chain called in individual stages as well as for the overlap groups, yielding the signal in each stage relative to all peaks called by one or more samples. A final level of filtering retains only those peak regions with a maximum RPKM in any stage of 2.0 and a difference from the lowest to the highest stage RPKM of at least 0.75 times the maximum stage value.

##### Peak cluster/trajectory analysis

To assess whether peak regions formed distinct clusters or conformed better to a continuous trajectory of types, we used the umap2 function from the R uwot package to perform various types of UMAP analysis of peak region values for each stage. One UMAP type used stage RPKM values and Euclidean distances and thus took ATAC-seq magnitude into account. For other analyses, region RPKM scores were scaled as the log2 fold change from the mean RPKM and analyzed using both Euclidean and correlation distances. All outputs were consistent with a continuous trajectory of peak region types that correlated well with stage mean.

##### Peak lag analysis

We performed lag analysis to understand how peak regions correlated to each other when they co-localized in the genome. We calculated scaled log2 fold RPKM values as above and calculated the degree of variance between peak regions as a function of the binned log10 distance between them. Lower variance at lower inter-peak distances reveals the degree of auto-correlation of nearby peaks and the distance over which it manifests.

##### Re-analyzing bins without dinucleosome peaks

A possible confounder of the 1 kb bin analyses above is that bins overlapping hyper-accessible peaks might behave differently than the rest of the genome. To assess the magnitude of this effect we used pipeline action ‘atac recollatè to repeat the bin-level analyses after masking peak-containing bins as “not included”. Doing so had little impact on binned results because there are many more non-peak than peak-containing bins in the genome.

##### Code availability

The protamine-tools pipeline is available on GitHub at https://github.com/HammoudWilson/protamine-tools. Within the repository, (i) the ‘templates’ folder contains example job files to support application of the pipeline code to other similar data sets, (ii) the ‘resources’ folder contains a small files needed to run the pipeline, including the compiled Rust executable for ab_initio peak calling, and (iii) the ‘logs’ folder contains files with complete job file configuration and logged output from jobs as we ran them. No additional installed programs are required to run the protamine-tools pipeline as it contains commands to build a conda environment with appropriate program versions.

## Comparison between peak calls from pipeline 1 and pipeline 2

The number of peaks called from methods discussed above were plotted as bar plots using ggplot2 geom_bar function in R. Overlapping peaks between the two methods were identified using the findoverlaps function from the package GenomicRanges and visualized as Venn diagrams plotted using the R package called VennDigram. Peaks were assigned to different genomic regions using the annotatePeak function from the package ChIPseeker in R.

## ATAC MACS2 and Di-nucleosome Heatmap

To visualize temporal chromatin accessibility dynamics, a union peakset was generated by merging MACS2 narrowPeak files across seven developmental stages (early RS to late ES) and reducing overlapping regions using the GenomicRanges^20^ (v1.58.0) R package. GC-residual z-score signal intensities were extracted by overlapping these union peaks with previously mentioned 1kb genomic bins and calculating the mean signal per peak. Following signal centering, the dataset was filtered to include only dynamic regions, defined as peaks with a signal range (max – min) greater than 1.0 across all stages. For visualization, a representative subset of 50,000 peaks was ordered by their stage mean. The final signal matrix was row-standardized using z-score normalization and plotted using the pheatmap (v1.0.13)^21^ R package, with the color scale truncated at ±3 to emphasize relative accessibility shifts across the developmental trajectory.

To visualize chromatin accessibility dynamics using dinucleosome peak calls, adjacent peaks that overlap in their nucleosomes or flanking nucleosome-free regions are merged into a “nucleosome chain”, and gap lengths weighted by the peak scores are referred to as stage mean. The combined matrix was ordered by stage mean (calculation described in earlier section) and plotted using pheatmap (v1.0.13)^21^ R package

## Processing CUT&Tag data

Cut&Tag reads were aligned using Novoalign (version 4.04.01) to a combined mouse (UCSC version mm39) and *Drosphila* (UCSC version BDGP6) reference genome using default-tuned parameters for NOVASEQ and including built-in adapter trimming with sequence CTGTCTCTTATACACATCT. Alignments were tagged with mate scores using Samtools fixmate (version 1.21). Alignments were split into separate genome-specific files using Picard ReorderSam (version 2.23.3).

Alignment files from samples and replicates were analyzed using the MultiRepMacsChIPSeq pipeline (https://huntsmancancerinstitute.github.io/MultiRepMacsChIPSeq/, release 20). This included fragment filtering based on insertion length (40-300 bp), proper pairs, exclusion of mitochondrial and contig chromosomes, removal of optical duplicate alignments (pixel distance of 250), and partial PCR de-duplication via random subsampling to a final total duplication rate of 5%.

The fragment coverage bigWig tracks were generated by the MultiRepChIPSeq pipeline after filtering alignments for proper pairs, mapping quality (> 10), insert size (40-300 bp), exclusion from unwanted chromosomes (chrM and unmapped contigs) and exclusion intervals (extreme coverage over repetitive sequence elements), and fragment de-duplication (removal of all optical duplicates and random subsampling of remaining biological/PCR duplicates to 5 or 10% of non-duplicate levels. The tracks were scaled to Reads (Fragments) Per Million (RPM).

The pipeline was run with the --independent flag to generate independent peak calls of each sample replicate. Replicate peak calls were then merged by the pipeline, retaining only those intervals called by 2 or more replicates as the final sample peak call set.

## Hi-C Data Processing and Analysis

### Data Processing and Quality Control

Raw sequencing data, compressed in ORA format, were decompressed using Orad (v2.7.0-c9253; Python 3.11) referencing the NCBI RefSeq assembly GCF_000001635.27. Subsequent processing was performed using a custom implementation of the Juicer pipeline^22^(v1.6) adapted for a cluster environment. Reads were aligned to the mm39 reference genome (ENCODE no-alt analysis set) using BWA-MEM (v0.7.17). Ligation contacts were validated against restriction sites for Arima-HiC (motifs GATC and GANTC). PCR duplicates and reads with a MAPQ score < 30 were removed to retain only valid contact pairs. Hi-C contact matrices were generated using Juicer Tools (v1.7.6; OpenJDK 1.8.0_452) and balanced using Knight-Ruiz (KR) normalization. Replicate files were merged using the Juicer mega.sh script to increase read depth for downstream analysis. The HiCRep reproducibility measure was run using 3DChromatin_ReplicateQC^23,24^ v0.0.1 with parameters ‘h=5, maxdist=5000000’ at 50 kb Hi-C resolution.

### Data Analysis

Compartments (A/B) were identified using the Juicer Tools eigenvector command at 250kb resolution. To ensure consistency across samples, the sign of the eigenvector was oriented such that positive values corresponded to the A compartment in the ES-E14 mESC cell line^25^. This orientation and subsequent comparative plotting were performed using custom Python scripts (Python 3.11, pandas 2.3.1, NumPy 2.3.1, Matplotlib 3.10.3, SciPy 1.16.0). Genomic coordinates were converted from mm10 to mm39 using the UCSC liftOver tool with the mm10ToMm39.over.chain.gz chain file. 2D pairwise contacts of Hi-C data were visualized with Juicebox (v2.15)^26^ at a 500 kb resolution with a coverage normalization, unless specified otherwise.

### Assessing ATAC-seq Peaks across Stages

For each stage, we sought to assess the significance of the number of MACS2 peaks per chromosome. We constructed a null distribution by repeatedly sampling with replacement (1,000 permutations) peak files at random (numpy v2.3.1) for each chromosome and summing their counts, ensuring equal representation across stages. Observed counts were then compared to this null distribution to calculate z-scores and p-values (scipy v1.16.0), which were adjusted for multiple testing using the Benjamini-Hochberg^7^ method with FDR of 0.05 (statsmodels v0.14.5). To account for correlation between stages, pairwise Jaccard indices^8,9^ (pybedtools v0.12.0) between peak sets were used as proxies for correlation and we applied Brown’s method^10^, a correlation-corrected extension of Fisher’s combined probability test^11^, to evaluate the global deviation of observed data from the null. The covariance terms required for Brown’s method were estimated using the Kost & McDermott polynomial approximation^12^. Results were visualized as boxplots (matplotlib v3.10.3, seaborn v0.13.2) of the null distributions with observed values and statistical significance annotated. The global p-values indicate deviance from an expected random background if there were not differences between stages. The analysis above was conducted in a python (v3.11.13) notebook using pandas v2.3.1.

### Linear Mixed Model Regression Analyses Data Processing

We integrated transcription and chromatin accessibility data for the linear mixed model regression analysis. Specifically, scRNA-seq transcripts were annotated using the *Mus musculus* genome assembly (GRCm38.81). To remove gene duplicates, we kept the sources in order ensembl_havana, havana, then ensembl. If there were still duplicates, we kept the lower gene_version. We utilized PyRanges^13^ (v0.1.4) join to intersect transcription data with ATAC-seq GC-corrected residuals. For each gene, chromatin accessibility was calculated as the mean signal at the bin containing the promoter the bins upstream and downstream of the promoter. Data was formatted into a longitudinal structure where each gene-stage observation constituted a single data point. The analysis above was conducted in python (v3.11.13).

### Quantifying Transcription-Accessibility Coupling Strength (Fig 5a)

#### Data Preprocessing

Genes encoding protamines (*Prm1, Prm2*) and transition proteins (*Tnp1, Tnp2*) are hyper-upregulated during sperm packaging in the early ES stage from driving global regression trends and were resultingly removed from subsequent analysis. Additionally, inactive genes with mean expression across stages below the 25th percentile were excluded.

#### Statistical Modeling

We fitted a Linear Mixed-Effects Model (LMM) using the lme4 package^14^ (v1.1.38) in R (v4.5.2). To assess the relationship between transcription and chromatin accessibility across stages, the model was specified as follows:

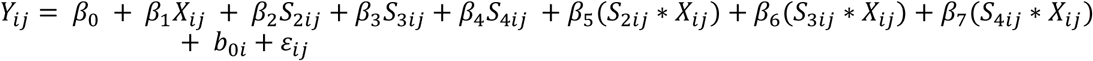

where

● 𝑌_𝑖𝑗_ is the ATAC GC residual corrected Z-score for gene 𝑖 at stage 𝑗.
● 𝑋_𝑖𝑗_ is the log(Transcription + 1) for gene 𝑖 at stage 𝑗.
● 𝑆_2𝑖𝑗_, 𝑆_3𝑖𝑗_, 𝑆_4𝑖𝑗_: Binary indicator variables taking the value 1 if observation *j* for gene *i* corresponds to stages int RS, late RS, or early ES, respectively, and 0 otherwise. (Reference stage: early RS).
● 𝑏_0𝑖_ is the random intercept for gene *i*, accounting for baseline heterogeneity.
● 𝛽_1_: The slope of the transcription-accessibility relationship for the reference stage.
● 𝛽 _5−7_ : The interaction coefficients representing the difference in the transcription slope for stages int RS, late RS, and early ES relative to the reference stage.
● 𝜖_𝑖𝑗_ : The residual error for gene 𝑖 at stage 𝑗, assumed 𝑁(0, 𝜎_𝜖_ ^2^)

#### Significance Testing and Diagnostics

P-values for fixed effects were obtained using Satterthwaite’s degrees of freedom method via the lmerTest package^15^. Model fit was assessed using residual plots and QQ-plots. Visual inspection indicated minor heteroscedasticity; however, we decided to continue with the model because parameter estimates in LMMs remain unbiased even under heteroscedasticity, though precision may be affected^16^.

#### Data Visualization

To dissect the interaction effects, we used the emmeans package (v2.0.1, https://rvlenth.github.io/emmeans/). Specifically, we used emtrends to estimate the slope of the transcription-accessibility relationship and its asymptotic 95% confidence intervals for each stage. Subsequently, we performed pairwise contrasts of these slopes to explicitly test for significant differences in coupling strength (Δβ) between all possible stage pairs, corrected for multiple comparisons using the Tukey method^18^. We also used emmip function to calculate marginal predicted ATAC values across the range of transcription levels.

Two types of visualizations were generated using ggplot2 (v4.0.1):

1. Marginal Prediction Plots: Fitted regression lines with confidence ribbons were plotted to visualize the global trend of chromatin coupling across stages.
2. Slope Comparison Plots: Point-range plots were generated to directly compare the estimated slopes (β) and their precision across stages.

#### Low vs High Expression Regression (Fig 5b)

All statistical analyses were conducted in R (v4.5.2). To analyze accessibility trajectories, genes were stratified based on their transcriptional activity in the early RS stage. “High” transcription genes were defined as the top 5th percentile, while “Low” genes were defined as those in the 25th–50th percentile range using dplyr (v1.1.4).

To control for initial differences in chromatin openness, we calculated the change in accessibility (Δ) from baseline for every gene at every timepoint (𝑌_𝑖𝑗_ − 𝑌_𝑖0_). We employed a LMM to characterize the differential rate of remodeling over time, accounting for the developmental inflection point at the transition from Round Spermatids to Elongating Spermatids. Developmental stages were mapped to a continuous time variable t (Early RS = 0, Int RS = 1, Late RS = 2, Early ES = 3). A knot was placed at t=2 to allow for distinct rates of change before and after the onset of elongation.

The model was specified as follows:

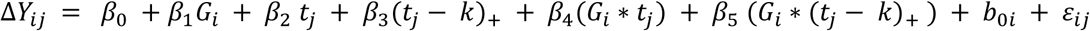

where

● Δ𝑌_𝑖𝑗_ is the change in ATAC Z-score relative to the Early RS baseline for gene 𝑖 at stage 𝑗.
● 𝐺_𝑖_ is the transcription group indicator (1 for High, 0 for Low).
● 𝑡_𝑗_ is the time point.
● (𝑡_𝑗_ − 𝑘)_+_ represents the linear spline function (0 before the knot k=2, and t−2 after the knot).
● 𝛽_4_and 𝛽_5_ capture difference in remodeling rate (velocity) between transcription groups before and after the knot, respectively.
● 𝑏_0𝑖_ is the random intercept for gene 𝑖, accounting for gene-specific deviations from the group trajectory.

The model was fitted using the lme4 package^14^ (v1.1.38). We tested the null hypothesis H_0_: 𝛽_4_ = 𝛽_5_ = 0, interpreted as there is no difference in the rate of remodeling between low and highly transcribed genes using the car package’s^19^ (v3.1.3) linearHypothesis to calculate the Wald test statistic.

##### Data Visualization

Model-predicted means were generated for each group across all time points. 95% Confidence Intervals were calculated using the standard errors derived from the fixed-effects variance-covariance matrix. Visualization was performed using ggplot2 (v4.0.1).

#### Histone Modifications by Compartments **Supp Figure 5c/d**

A python (v3.9.23) script was used to annotate peak regions with compartment (A/B) and extract signal values from bedGraph files using pyBedGraph^20^ (v0.5.43). For each chromosome, the mean ATAC signal across each peak was computed in parallel for all peaks, and each region was assigned to a compartment based on the sign of the compartment score. The resulting annotated dataframes were then used for downstream statistical analysis and visualization of signal distributions by compartment. For each stage, we performed a two-sided Kolmogorov-Smirnov test^21^ (scipy v1.13.1) to assess differences between compartments, displaying the results on boxplots and Empirical Cumulative Distribution Function plots with annotated sample sizes and p-values using seaborn (v0.13.2) and matplotlib (v3.9.4).

#### Median ATAC-seq signal trace across stages

In a python notebook, we converted the ATAC .rds file to a pandas (v2.3.1) dataframe using the rpy2 (v3.6.3) python package. Each ATAC bin was matched to its corresponding compartment using genomic coordinates. For each stage and compartment, the median GC residual z-score was calculated.

To establish a baseline for comparison, a theoretical null distribution was generated by sampling 1,000 points from a standard normal distribution *N*(0,1), using a fixed random seed (42) from NumPy (v2.3.1) for reproducibility. The choice of *N*(0,1) as the background distribution is predicated on the properties of the GC residual z-scores; by definition, z-scores are standardized transformations where the global population mean is 0 and the standard deviation is 1.

Under the null hypothesis - assuming no systematic enrichment or depletion of ATAC signal within specific genomic compartments - the GC residuals should remain centered around the expected mean of 0. By comparing the observed compartment-specific medians against this theoretical expectation, we can identify biological deviations from the global genomic background. To ensure a fair comparison and account for sampling noise in the null model, we applied the same bootstrapping procedure (1,000 resamples, n=1,000) to both the synthetic and observed data. This involved subsampling 1,000 points from the observed data to match the size of the expected dataset, providing a robust estimate of the median and its associated 95% confidence intervals. The resulting confidence intervals and medians were visualized using matplotlib (v3.10.3).

## Data Availability

All sequencing data generated in this study have been deposited in the NCBI Sequence Read Archive (SRA) under BioProject accession number **PRJNA1432264**. The records are currently undergoing curator review and will be made publicly available upon completion of processing.

## Notes

### Competing Interest Statement

The authors have declared no competing interest.

