## Supplementary material for "3D chromatin compartment of round spermatids encodes the spatiotemporal program of histone-to-protamine exchange in spermiogenesis": Methods

**Catalog no of antibodies used:**

| **Antibodies** | **Catalog No:** |
| --- | --- |
| V5 (mouse) | MCA1360 |
| V5 (rabbit) | V8137 |
| Hup1n | MAb-Hup1N-150 |
| Hup2b | MAb-Hup2B-150 |
| Tnp1 | 17178-1-AP |
| Tnp2 | sc-393843 |
| H3 | ab1791 |
| H4ac | 06-866 |
| H3k27me3 | 13-0055 |
| H2B | 07-371 |
| H4 | 16047-1-AP |

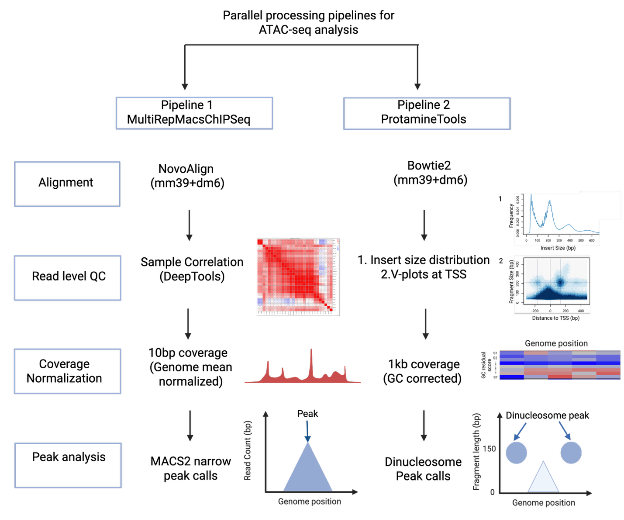

Parallel processing pipelines used for ATAC-seq analysis.

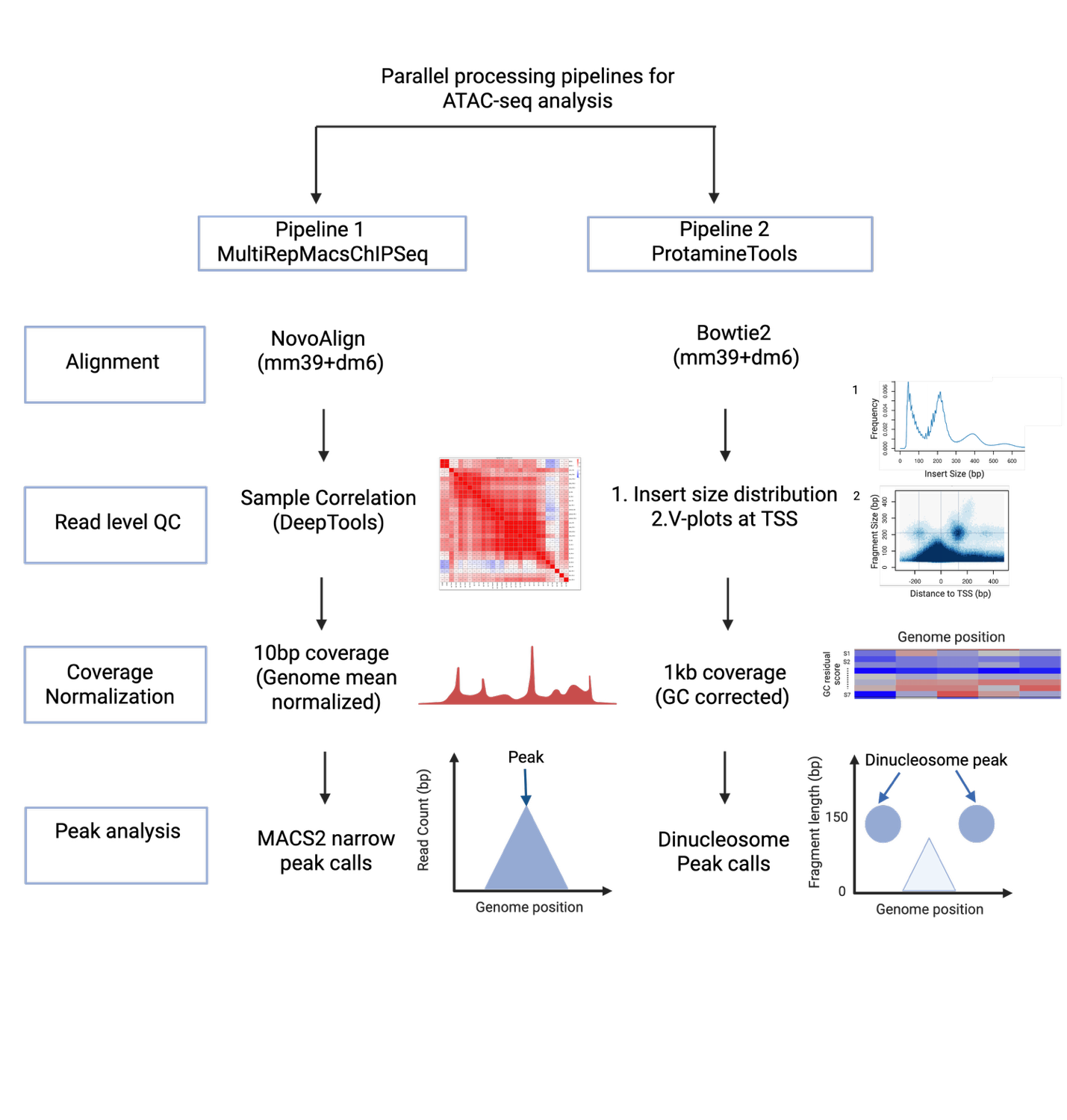

**ATAC-seq analysis – Pipeline1: MultiRepMacsChIPSeq**

Alignment and Normalization

ATAC-seq reads were aligned using Novoalign (version 4.04.01, http://novocraft.com) to a combined mouse (UCSC version mm39) and *Drosphila* (UCSC version BDGP6) reference genome using default-tuned parameters for NOVASEQ and including built-in adapter trimming with sequence CTGTCTCTTATACACATCT. Alignments were tagged with mate scores using Samtools fixmate (version 1.21, https://www.htslib.org/doc/1.21/samtools-fixmate.html). Alignments were split into separate genome-specific files using Picard ReorderSam (version 2.23.3, https://broadinstitute.github.io/picard/). Alignment files from samples and replicates were put through the MultiRepMacsChIPSeq pipeline (<https://huntsmancancerinstitute.github.io/MultiRepMacsChIPSeq/>, release 20) for fragment-based peak calling. Properly-paired alignments were filtered based on insertion length (40-300 bp), exclusion of mitochondrial and contig chromosomes, removal of optical duplicate alignments (pixel distance of 250), and partial PCR de-duplication via random subsampling to a final total duplication rate of 5%. Genome-wide fragment coverage was generated (in 10 bp bins), normalized to a depth of 1 million fragments. Coverage was scaled to the median sequencing depth of the given samples, statistical enrichment (q-value) genome-wide tracks generated using MACS2^11^ and the empirical genomic mean coverage, and peaks called using MACS2^11^ using a minimum peak size of 200 bp, gap of 50 bp, and minimum threshold of 0.01 (1% FDR).

$$Y_{ij}= \beta_{0} + \beta_{1}X_{ij} + \beta_{2}S_{2ij}+\beta_{3}S_{3ij}+\beta_{4}S_{4ij} +\beta_{5}{(S}_{2ij}*X_{ij})+\beta_{6}{(S}_{3ij}*X_{ij})+\beta_{7}{(S}_{4ij}*X_{ij}) + b_{0i}+\varepsilon_{ij}$$

where

- $Y_{ij}$ is the ATAC GC residual corrected Z-score for gene $i$ at stage $j$.
- $X_{ij}$ is the log(Transcription + 1) for gene $i$ at stage $j$.
- $S_{2ij}, S_{3ij}, S_{4ij}$: Binary indicator variables taking the value 1 if observation *j* for gene *i* corresponds to stages int RS, late RS, or early ES, respectively, and 0 otherwise. (Reference stage: early RS).
- $b_{0i}$ is the random intercept for gene *i*, accounting for baseline heterogeneity.
- $\beta_{1}$: The slope of the transcription-accessibility relationship for the reference stage.
- $\beta{}_{5-7}$: The interaction coefficients representing the difference in the transcription slope for stages int RS, late RS, and early ES relative to the reference stage.
- $\epsilon_{ij}$: The residual error for gene $i$ at stage $j$, assumed $N(0, {\sigma_{\epsilon}}^{2})$

The model was specified as follows:

$${\Delta Y}_{ij} = \beta_{0} +\beta_{1}G_{i} + \beta_{2} t_{j} + \beta_{3}(t_{j}- k)_{+} + \beta_{4}(G_{i}*t_{j}) + \beta_{5} (G_{i}*(t_{j}- k)_{+} ) + b_{0i} + \varepsilon_{ij}$$

where

- ${\Delta Y}_{ij}$ is the change in ATAC Z-score relative to the Early RS baseline for gene $i$ at stage $j$.
- $G_{i}$ is the transcription group indicator (1 for High, 0 for Low).
- $t_{j}$ is the time point.
- $(t_{j}- k)_{+}$ represents the linear spline function (0 before the knot k=2, and t−2 after the knot).
- $\beta_{4}$and $\beta_{5}$ capture difference in remodeling rate (velocity) between transcription groups before and after the knot, respectively.
- $b_{0i}$ is the random intercept for gene $i$, accounting for gene-specific deviations from the group trajectory.

1	Pepin, A. S., Jazwiec, P. A., Dumeaux, V., Sloboda, D. M. & Kimmins, S. Determining the effects of paternal obesity on sperm chromatin at histone H3 lysine 4 tri-methylation in relation to the placental transcriptome and cellular composition. *Elife* **13** (2024). <https://doi.org/10.7554/eLife.83288>

2	Moritz, L. *et al.* Sperm chromatin structure and reproductive fitness are altered by substitution of a single amino acid in mouse protamine 1. *Nat Struct Mol Biol* **30**, 1077-1091 (2023). <https://doi.org/10.1038/s41594-023-01033-4>

3	Nakata, H., Wakayama, T., Takai, Y. & Iseki, S. Quantitative analysis of the cellular composition in seminiferous tubules in normal and genetically modified infertile mice. *J Histochem Cytochem* **63**, 99-113 (2015). <https://doi.org/10.1369/0022155414562045>

4	Hogarth, C. A. *et al.* Turning a spermatogenic wave into a tsunami: synchronizing murine spermatogenesis using WIN 18,446. *Biol Reprod* **88**, 40 (2013). <https://doi.org/10.1095/biolreprod.112.105346>

5	Herrmann, C., Avgousti, D. C. & Weitzman, M. D. Differential Salt Fractionation of Nuclei to Analyze Chromatin-associated Proteins from Cultured Mammalian Cells. *Bio Protoc* **7** (2017). <https://doi.org/10.21769/BioProtoc.2175>

6	Green, C. D. *et al.* A Comprehensive Roadmap of Murine Spermatogenesis Defined by Single-Cell RNA-Seq. *Dev Cell* **46**, 651-667 e610 (2018). <https://doi.org/10.1016/j.devcel.2018.07.025>

7	Shen, Y. C. *et al.* TCF21(+) mesenchymal cells contribute to testis somatic cell development, homeostasis, and regeneration in mice. *Nat Commun* **12**, 3876 (2021). <https://doi.org/10.1038/s41467-021-24130-8>

8	Gill, M. E., Kohler, H. & Peters, A. Dual DNA staining enables isolation of multiple sub-types of post-replicative mouse male germ cells. *Cytometry A* **101**, 529-536 (2022). <https://doi.org/10.1002/cyto.a.24539>

9	Corces, M. R. *et al.* An improved ATAC-seq protocol reduces background and enables interrogation of frozen tissues. *Nat Methods* **14**, 959-962 (2017). <https://doi.org/10.1038/nmeth.4396>

10	Kaya-Okur, H. S. *et al.* CUT&Tag for efficient epigenomic profiling of small samples and single cells. *Nat Commun* **10**, 1930 (2019). <https://doi.org/10.1038/s41467-019-09982-5>

11	Zhang, Y. *et al.* Model-based analysis of ChIP-Seq (MACS). *Genome Biol* **9**, R137 (2008). <https://doi.org/10.1186/gb-2008-9-9-r137>

12	Ramirez, F., Dundar, F., Diehl, S., Gruning, B. A. & Manke, T. deepTools: a flexible platform for exploring deep-sequencing data. *Nucleic Acids Res* **42**, W187-191 (2014). <https://doi.org/10.1093/nar/gku365>

13	Quinlan, A. R. & Hall, I. M. BEDTools: a flexible suite of utilities for comparing genomic features. *Bioinformatics* **26**, 841-842 (2010). <https://doi.org/10.1093/bioinformatics/btq033>

14	Huang, D. W. *et al.* DAVID Bioinformatics Resources: expanded annotation database and novel algorithms to better extract biology from large gene lists. *Nucleic Acids Res* **35**, W169-175 (2007). <https://doi.org/10.1093/nar/gkm415>

15	Chen, S. fastp 1.0: An ultra-fast all-round tool for FASTQ data quality control and preprocessing. *Imeta* **4**, e70078 (2025). <https://doi.org/10.1002/imt2.70078>

16	Amemiya, H. M., Kundaje, A. & Boyle, A. P. The ENCODE Blacklist: Identification of Problematic Regions of the Genome. *Sci Rep* **9**, 9354 (2019). <https://doi.org/10.1038/s41598-019-45839-z>

17	Pockrandt, C., Alzamel, M., Iliopoulos, C. S. & Reinert, K. GenMap: ultra-fast computation of genome mappability. *Bioinformatics* **36**, 3687-3692 (2020). <https://doi.org/10.1093/bioinformatics/btaa222>

18	Green, B., Bouchier, C., Fairhead, C., Craig, N. L. & Cormack, B. P. Insertion site preference of Mu, Tn5, and Tn7 transposons. *Mob DNA* **3**, 3 (2012). <https://doi.org/10.1186/1759-8753-3-3>

19	Kaye, E. G. *et al.* RNA polymerase II pausing is essential during spermatogenesis for appropriate gene expression and completion of meiosis. *Nat Commun* **15**, 848 (2024). <https://doi.org/10.1038/s41467-024-45177-3>

20	Lawrence, M. *et al.* Software for computing and annotating genomic ranges. *PLoS Comput Biol* **9**, e1003118 (2013). <https://doi.org/10.1371/journal.pcbi.1003118>

21	Kolde, R. *pheatmap: Pretty Heatmaps*, <<https://github.com/raivokolde/pheatmap>> (2025).

22	Durand, N. C. *et al.* Juicer Provides a One-Click System for Analyzing Loop-Resolution Hi-C Experiments. *Cell Syst* **3**, 95-98 (2016). <https://doi.org/10.1016/j.cels.2016.07.002>

23	Yang, T. *et al.* HiCRep: assessing the reproducibility of Hi-C data using a stratum-adjusted correlation coefficient. *Genome Res* **27**, 1939-1949 (2017). <https://doi.org/10.1101/gr.220640.117>

24	Yardimci, G. G. *et al.* Measuring the reproducibility and quality of Hi-C data. *Genome Biol* **20**, 57 (2019). <https://doi.org/10.1186/s13059-019-1658-7>

25	Wang, X. *et al.* A generalizable Hi-C foundation model for chromatin architecture, single-cell and multi-omics analysis across species. *bioRxiv* (2024). <https://doi.org/10.1101/2024.12.16.628821>

26	Durand, N. C. *et al.* Juicebox Provides a Visualization System for Hi-C Contact Maps with Unlimited Zoom. *Cell Syst* **3**, 99-101 (2016). <https://doi.org/10.1016/j.cels.2015.07.012>
