## Supplementary Figures for "3D chromatin compartment of round spermatids encodes the spatiotemporal program of histone-to-protamine exchange in spermiogenesis"

#### Supplementary Figure 1

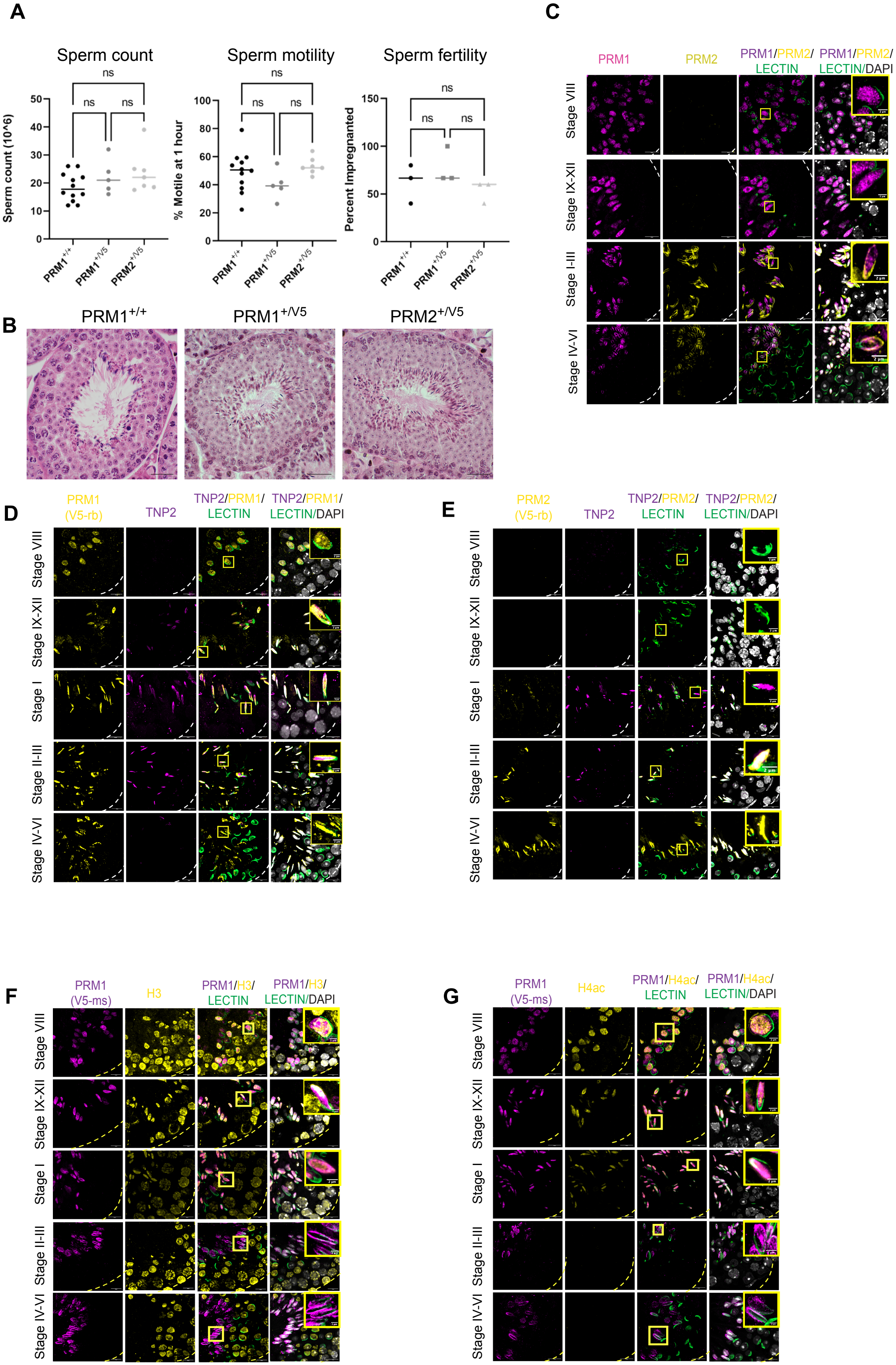

### Supplementary Figure 2

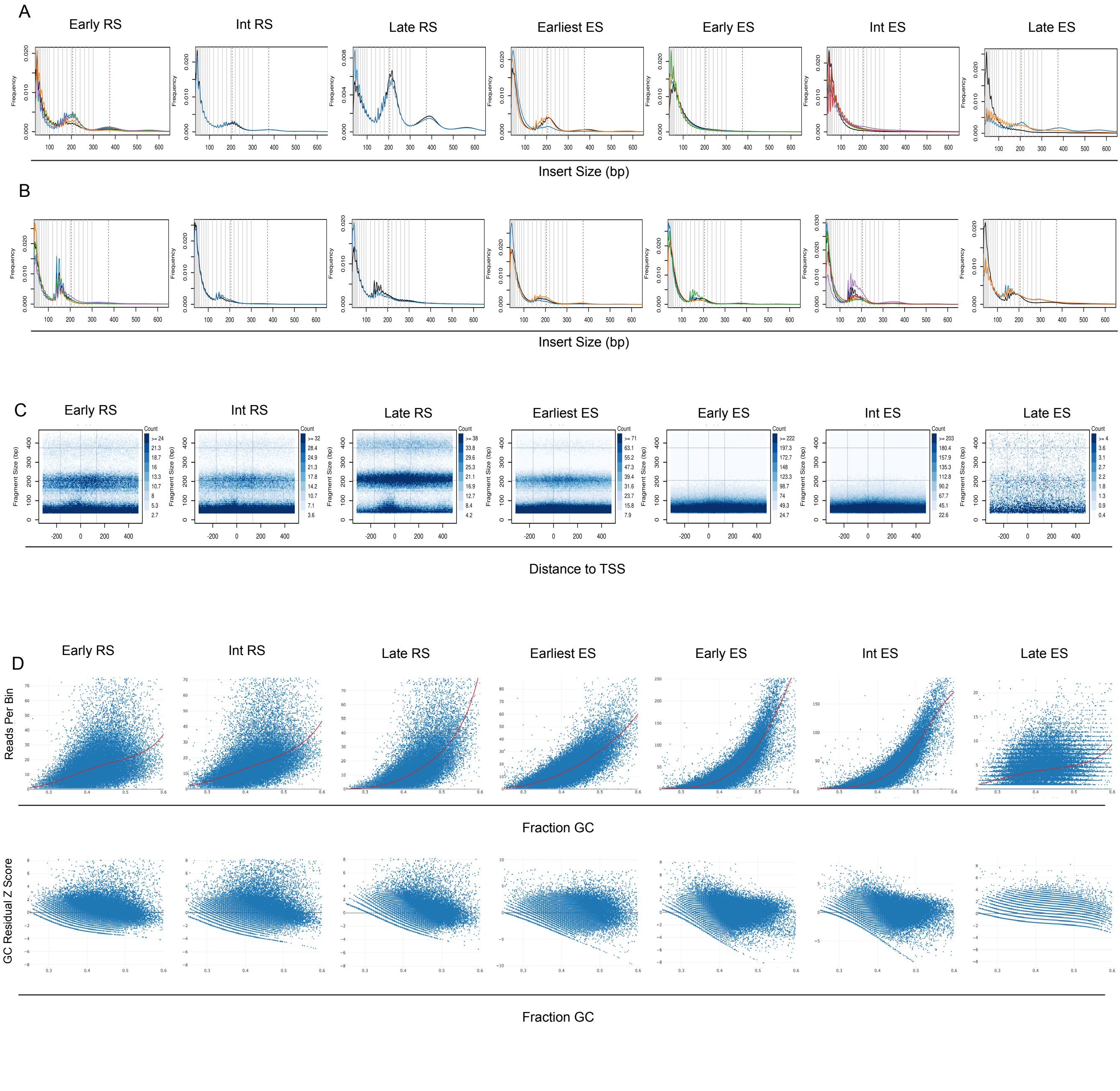

### Supplementary Figure 3

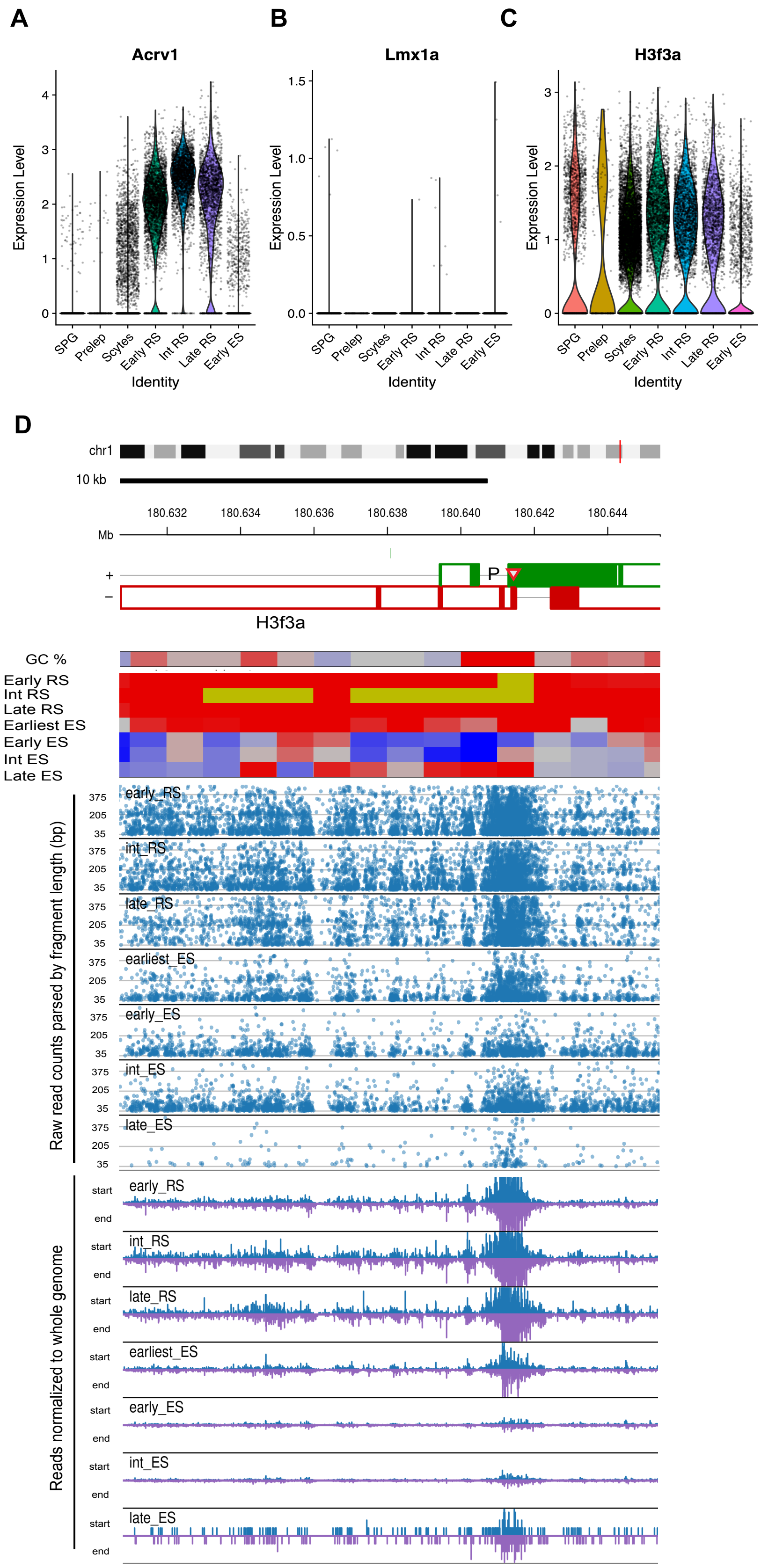

### Supplementary Figure 4

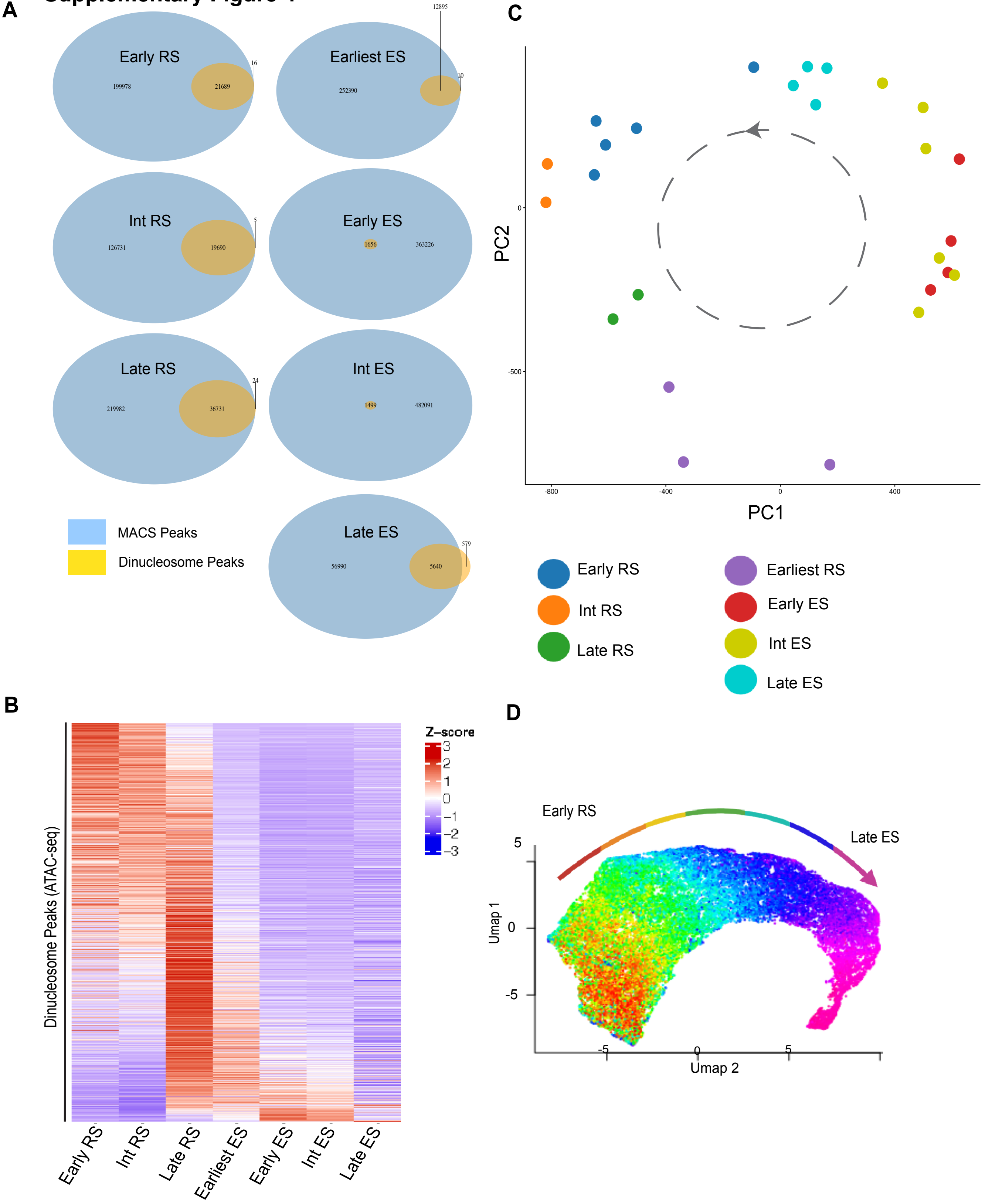

### Supplementary Figure 5

**A**

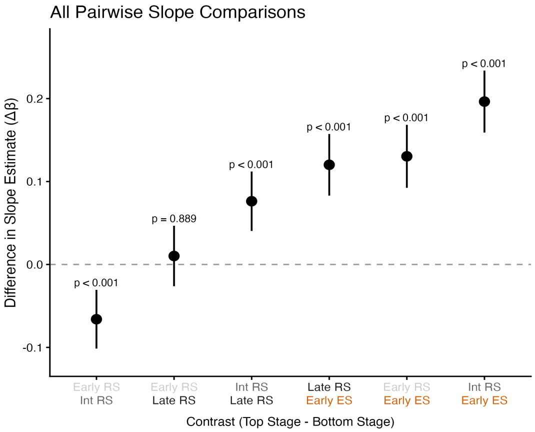

### Supplementary Figure 6

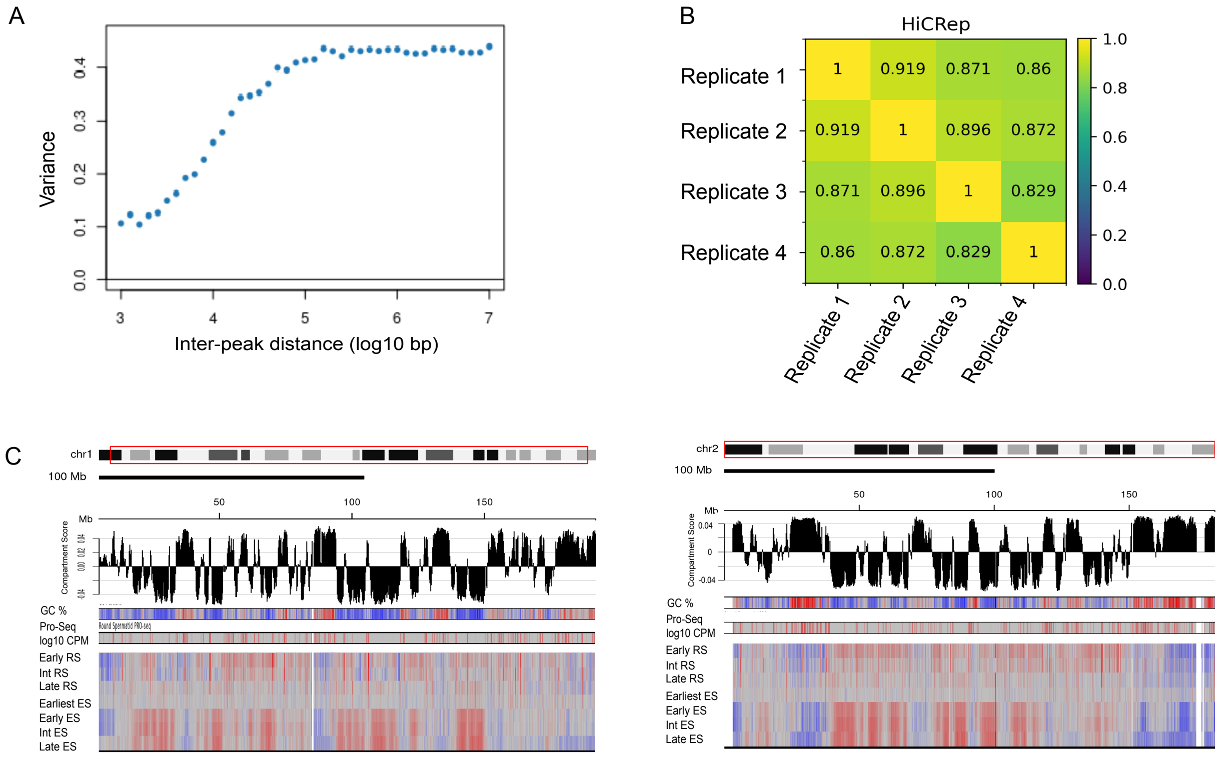

### Supplementary Figure 7

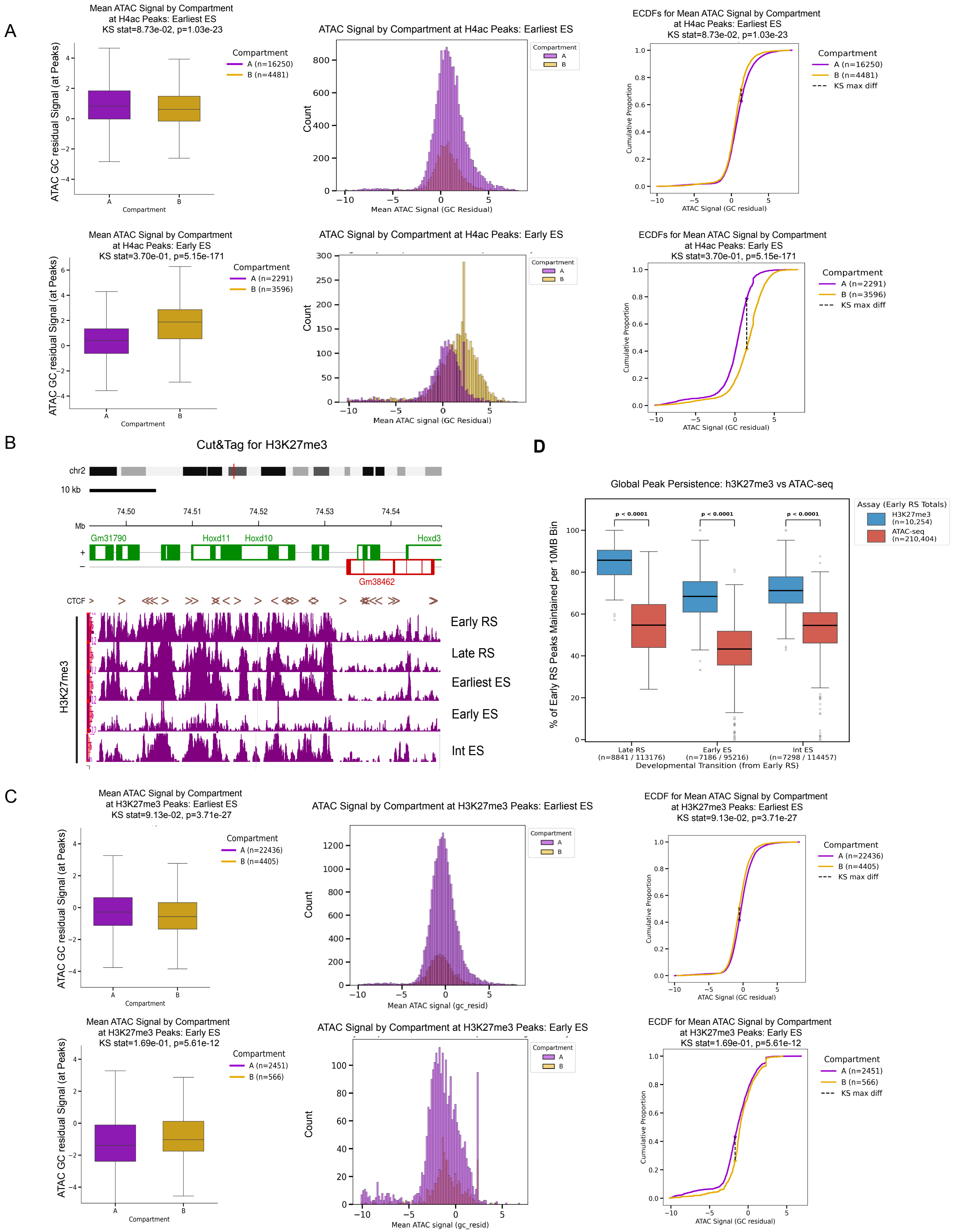
