## Supplementary Legends for "3D chromatin compartment of round spermatids encodes the spatiotemporal program of histone-to-protamine exchange in spermiogenesis"

**Supplementary Figure 1: Tagging the endogenous PRM1 and PRM2 loci has no effect on fertility or nuclear localization.**

**(A)** Prm1^⁺/V5^  and  Prm2^⁺/V5^ males have normal sperm counts, motility, and fertility relative to wild-type Prm1^⁺/⁺^ controls. Each data point represents an individual animal. Comparisons were made by one-way ANOVA with correction for multiple testing; ns, not significant.
**(B)** Hematoxylin and eosin staining of representative testis cross-sections from adult wild-type, Prm1^⁺/V5^ and Prm2^⁺/V5^ males. Scale bar, 20 µm.
**(C)** Testis cross-sections from Prm2^⁺/V5^ mice were co-stained with anti-V5 (PRM2) and Hup1N (PRM1). Magenta, PRM1 (Hup1N); yellow, PRM2 (V5); green, lectin; gray, DAPI. Scale bar, 10 µm; inset scale bar, 2 µm.
**(D)** Prm1^⁺/V5^ testis cross-sections were co-stained with anti-V5 (PRM1) and anti-TNP2. Magenta, PRM1 (V5); yellow, TNP2; green, lectin; gray, DAPI. Scale bar, 10 µm; inset scale bar, 2 µm.
**(E)** Prm2^⁺/V5^ testis cross-sections were co-stained with anti-V5 (PRM2) and anti-TNP2. Magenta, PRM2 (V5); yellow, TNP2; green, lectin; gray, DAPI. Scale bar, 10 µm; inset scale bar, 2 µm.
**(F,G)** Prm1^⁺/V5^ testis cross-sections were co-stained with anti-V5 (PRM1) and anti-H3 or anti-H4ac. Magenta, PRM1 (V5); yellow, H4ac; green, lectin; gray, DAPI. Scale bar, 10 µm; inset scale bar, 2 µm.

**Supplementary Figure 2: ATAC-seq replicates display concordant fragment length distributions.**
(**A)** ATAC-seq fragment length distributions for all biological replicates at each stage of spermiogenesis. Each line represents an individual replicate. Replicate numbers: Early RS, n = 5; Int RS, n = 2; Late RS, n = 2; Earliest ES, n = 3; Early ES, n = 5; Int ES, n = 6; Late ES, n = 3.
(**B)** Fragment length distributions of Drosophila spike-in reads corresponding to the samples in (A), used for normalization across stages.
(**C) V-plots of ATAC-seq fragment lengths across all seven stages as a function of distance from the annotated TSS of inactive genes based on PRO-seq (Kaye et al., 2024). Compared to active TSS in Fig. 2H, nucleosomes are not positioned but also disappear in Early ES and Int ES**.
(**D)** GC bias regression for ATAC-seq data across spermiogenesis stages. **(Top)** Reads per 1kb bin as a function of GC content. Red lines are the regression fit. **(Bottom)** Z-score relative to the GC regression fit as a function of GC content showing flattening of the data. Representative plots are shown for each stage.

**Supplementary Figure 3: Chromatin accessibility dynamics at loci with distinct functional properties during spermiogenesis are not affected by genome-wide normalization.**
**(A–C)** Single-cell RNA-seq expression profiles of Acrv1 (A), Lmx1a (B), and H3f3a (C) across spermatogenic cell types from mouse testis (Green et al., 2018). Cell centroids for each stage are indicated on the x-axis; expression level (normalized counts) on the y-axis.
**(D)** Chromatin accessibility at the H3f3a locus, representing a region that remains accessible throughout spermiogenesis with accessibility retained at the promoter in late elongating spermatids (late ES). **(Top)** Heatmap of GC-residual Z scores across the indicated spermatogenic stages. **(Middle)** V-plots across all seven stages. Each dot is a single mapped insert. Horizontal lines are at 35bp (minimum allowed fragment size), 205 bp (mono-nucleosome peak center), and 375 bp (di-nucleosome peak center). **(Bottom)** Strand-specific read counts normalized to the whole genome, enabling quantitative comparison of accessibility across stages independent of sequencing depth and providing strand-resolution of transcription factor footprints and precise nucleosome positioning at this locus.

**Supplementary Figure 4: Comparison of peak-calling strategies across spermiogenic stages.**
**(A)** Venn diagrams illustrating the overlap between peaks called by MACS2 (blue) and the dinucleosome peak-calling algorithm (gold) at each stage of spermiogenesis. Numbers indicate peak counts within each region.
**(B)** Heatmap of dinucleosome peak accessibility scores across spermiogenic stages. Each row is a peak overlap region ordered by the centroid of the stage-specific signals.
**(C)** Principal component analysis (PCA) of 26 ATAC-seq datasets using GC-residual scores computed over MACS2 peak regions. Each point represents an individual dataset, colored by spermatogenic stage as indicated. The samples cluster into a circular trajectory with Late ES returning to be most like Early RS.
**(D)** UMAP of di-nucleosome peaks across spermiogenesis, rainbow-colored by stage. Early RS (red), int RS (orange), late RS (yellow), earliest ES (green), early ES (cyan), int ES (blue) and Late ES (magenta). With (B), no discrete clusters are observed within a continuous trajectory of relative accessibility timing.

**Supplementary Figure 5: Comparing chromatin-transcription relationships during spermiogenesis.**
Forest plot displaying all pairwise contrasts of the transcription-accessibility slope estimates between stages. The y-axis represents the difference in slope for each comparison (Top Stage - Bottom Stage). Error bars denote 95% confidence intervals corrected for multiple comparisons (Tukey method). Comparisons involving Early ES consistently show a large difference, confirming that the coupling strength in this stage is significantly lower than in all RS stages. P-values indicate the statistical significance of the difference in slope.

**Supplementary Figure 6: Chromatin remodeling during spermiogenesis follows a reproducible, genome-wide, 3D-encoded program.**
**(A)** Variogram, i.e., lag analysis, depicting variance in GC-residual scores in ATAC-seq signal as a function of inter-peak distance (log10 bp).
**(B)** HiCRep stratum-adjusted correlation coefficients between all pairwise combinations of four Hi-C replicates.
(**C)** Genome-wide compartment scores and chromatin accessibility dynamics across chromosomes 1 (left) and 2 (right). From top to bottom: chromosome ideogram, compartment score track, GC content, Pro-Seq log10 CPM, and ATAC-seq GC-residual heatmaps across spermatogenic stages (Early RS through Late ES). B compartment (negative compartment scores) regions consistently correlate with higher relative accessibility in ES.

**Supplementary Figure 7: Compartment identity overrides histone acetylation in directing compaction order, while H3K27me3 acts as a stabilizing chromatin bookmark throughout spermiogenesis.
(A)** Boxplots (left), histograms (middle), and ECDFs (right) of mean ATAC-seq signal (GC residual) within H4ac peaks, stratified by A (purple) and B (gold) compartment identity, at earliest ES (top row) and early ES (bottom row).
**(B)** Cut&Tag signal for H3K27me3 at the Hoxd gene cluster on chromosome 2 across Early RS, Late RS, Earliest ES, Early ES, and Int ES stages. CTCF binding sites are indicated above the gene track.
**(C)** Boxplots (left), histograms (middle), and ECDFs (right) of mean ATAC-seq signal (GC residual) within H3K27me3 peaks, stratified by A (purple) and B (gold) compartment identity, at earliest ES (top row) and early ES (bottom row). KS statistics and p-values are indicated above each panel.
**(D)** Boxplots comparing the percentage of early RS peaks maintained per 10 Mb bin across three developmental transitions for H3K27me3 (blue) and ATAC-seq (red).
